# Exploiting threonine sidechains as molecular switches to modulate the fluorescence of genetically encoded biosensors

**DOI:** 10.1101/2022.04.25.489330

**Authors:** Lee Min Leong, Sang Chul Shin, Jun Kyu Rhee, Heejung Kim, Jihye Seong, Junhyuk Woo, Kyungreem Han, Douglas Storace, Bradley J. Baker

## Abstract

Rapid and reproducible optical transitions of a fluorescent protein (FP) can be achieved with a Genetically Encoded Voltage Indicator (GEVI) via manipulation of the membrane potential. These transitions revealed novel effects of internal mutations near the chromophore that would not be detected under steady state conditions. Mutating an internal threonine (T203) affected the speed of the voltage-dependent fluorescence transition suggesting a conformational change inside the protein. These optical transitions also demonstrated interplay between internal and externally oriented sidechains of the β-can structure. Replacing the steric hindrance of a phenylalanine near the chromophore with threonine (F165T) did not alter the resting fluorescence but resulted in a more complex fluorescent transition providing evidence for a flexible chromophore undergoing conformational changes. F165T orientation was influenced by the flanking external amino acids at positions 164 and 166 with 164F/165T/166T exacerbating the complexity of the voltage-dependent transition while 164T/165T/166F reduced the flexibility of the chromophore resembling the transition pattern of the original F165 version. Alphafold predictions reveal a threonine switch with different orientations of the F165T internal side chain depending on the direction of the offset in polarity at external positions 164 and 166. The crystal structures of the pH-sensitive FP, Super Ecliptic pHluorin and two derivatives solved in varying pH conditions also indicate interactions between the external protein surface and the internal environment providing another example of a threonine switch near the chromophore at T203. This ability to orient internal sidechains has led to the development of a novel GEVI that gets brighter upon depolarization of the plasma membrane, works at low light levels, is less susceptible to physiological pH, and provides in vivo signals. These observations affecting fluorescent transitions should also prove valuable to the development of any FP-based biosensor.

## Introduction

An ideal fluorescent protein (FP) exhibits bright and stable fluorescence. An ideal fluorescent biosensor has an additional requirement: multiple fluorescent states. The larger the change in photons (ΔF) between the different states, the better the signal-to-noise thereby facilitating in vivo recordings. The varying states of fluorescence for a biosensor must therefore also be stable so that ΔF is a reliable indicator of the change in the environmental condition being monitored. For instance, Super Ecliptic pHluorin (SE) is a pH sensitive FP that is dim at pH 5 [1]. Raise the pH incrementally and SE will get increasingly brighter until reaching a maximal fluorescence output around pH 8. However, when the pH is maintained, the fluorescence remains constant. Understanding how this environmental sensitivity is achieved would help improve the optical response of genetically encoded sensors.

FPs consist of an 11 β-sheet can structure that surrounds the internal chromophore. This organization protects the chromophore from the aqueous environment that would quench the fluorescence as well as provides support to minimize the flexibility of the chromophore. The internal environment surrounding the chromophore is therefore relatively stable consisting of hydrogen bond networks between the chromophore and the β-can structure as well as Van der Waal forces providing steric hindrance to prevent chromophore twisting. FP-based biosensors must therefore develop a mechanism for external environmental conditions to disrupt the internal environment of the chromophore.

The fluorescent properties of FPs are determined by the chemistry of the chromophore and the surrounding environment. For example, the protonated chromophore of avGFP is excited at 390 nm while the anionic form is excited at 470 nm [2-5]; yet both forms of the chromophore emit at nearly the same wavelength around 505 nm. The crystal structure of the FP revealed that the long Stokes shift for the protonated chromophore is due to a proton wire that allows rapid deprotonation of the chromophore upon excitation [6, 7]. While informative, crystal structures only provide a snapshot of stable conformations. Of further interest would be the rapid and reversible alteration of the internal environment of the FP in order to study the mechanism(s) inducing fluorescent transitions.

The genetically encoded voltage indicator (GEVI), ArcLight [8], is a biosensor that allows rapid transition of fluorescent states via control of the membrane potential of a cell. When the plasma membrane potential is held at −70 mV, the fluorescence is relatively bright. Depolarize the membrane and the fluorescence of ArcLight will dim. This consistent, rapid, and reversible change in fluorescence provides a valuable tool for exploring the effects amino acid substitutions have on transitioning to different fluorescent states.

ArcLight consists of a voltage-sensing domain (VSD) from the *Ciona intestinalis* voltage-sensing phosphatase fused to the pH-sensitive FP SE mentioned above (Figure 1A). The pH sensitivity of the FP domain is not sufficient for generating the large, voltage-dependent optical signal [8]. A mutation to the external surface of the β-can, SE A227D (numbering of amino acids throughout is based on the position in the FP), improves the voltage-dependent optical signal over 15-fold. Interestingly, the mutations rendering SE pH-sensitive also involve sidechains positioned on the outer surface of the β-can [1]. Pathways must therefore exist that convey external conditions to the internal chromophore creating an optical dependence on pH or, in the case of ArcLight with the A227D mutation, membrane potential.

**Figure 1.**
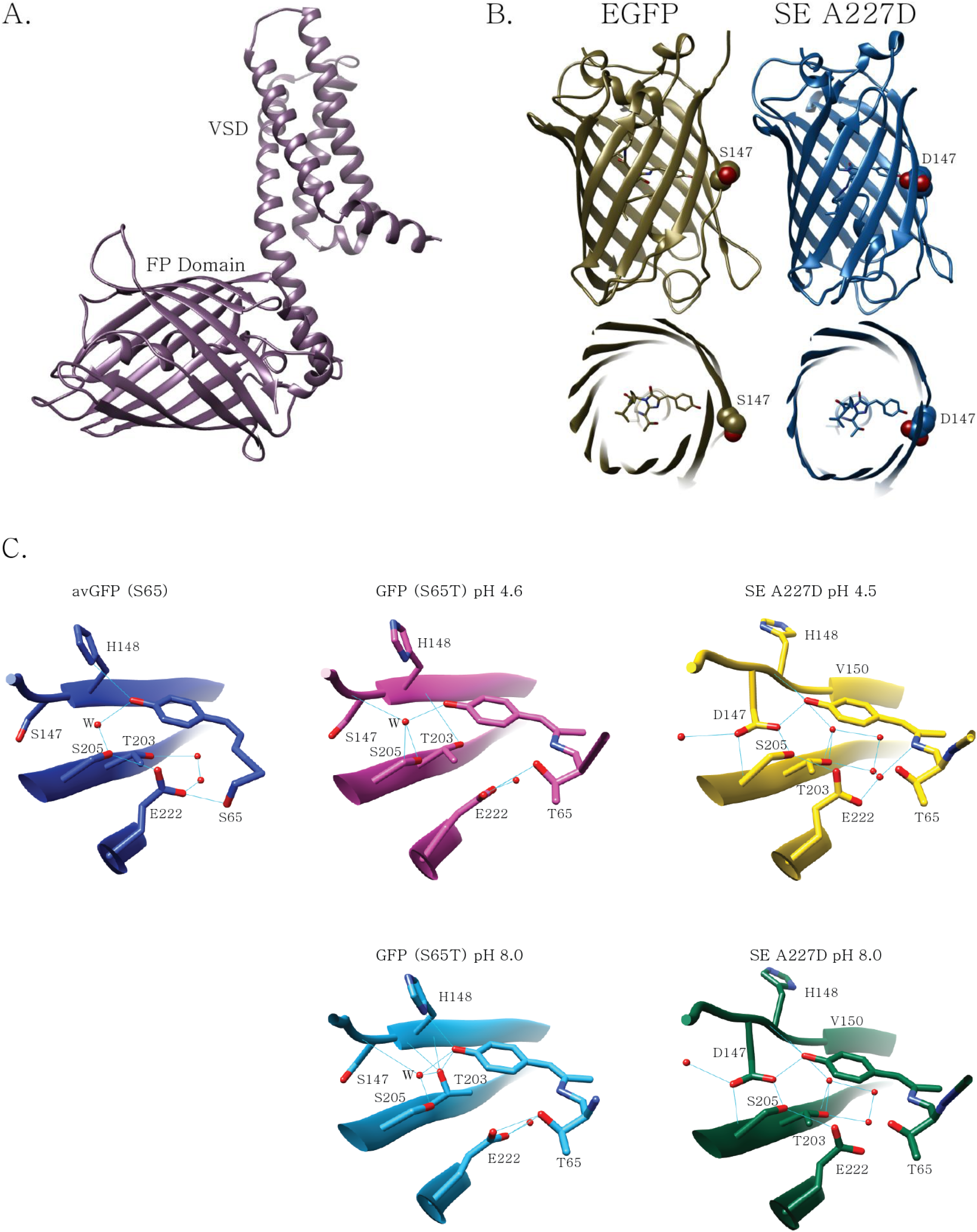
The crystal structure of SE A227D reveals a direct conduit between the chromophore and the external surface of the β-can. **A. Alphafold prediction of the ArcLight-derived GEVI, Triple Mutant (TM).** The four α-helices of the voltage-sensing domain (VSD) residing in the plasma membrane are fused to the fluorescent protein (FP) domain. The N-terminus random coil has been removed for clarity. **B. Comparison of the Super Ecliptic pHluorin A227D (SE A227D) crystal structure to EGFP.** The view from the side reveals the sidechain of the S147D mutation introduced during development of SE [1] has one oxygen exposed to the surface of the β-can structure while the other oxygen of the carboxylate sidechain forms a hydrogen bond with the chromophore. Oxygen molecules of S147 and D147 are shown in red. Top view demonstrates that the internal oxygen of the D147 sidechain is also internal providing a direct pathway from the external solvent to the chromophore. The SE A227D structure shown here was solved at pH 8.0. The structures of SE A227D and SE in various pH conditions are virtually identical (See supplemental figure S1). EGFP structure is shown on the left for comparison (PDB – 4EUL [12]). **C. Comparison of hydrogen bond networks involving the chromophores of avGFP, GFP (S65T), and SE A227D.** An internal view of the original avGFP (PDB – 1GFL [6]) is on the left showing the presence of a proton wire (hydrogen bonds are shown as light blue lines) from the chromophore to E222 via S205 and a water molecule (W).The structure of GFP (S65T) is also shown for two pH conditions (pH 4.6, PDB – 1C4F [13], and pH 8.0, PDB 1EMG [13]) to demonstrate how a change to the chromophore alters the orientation of E222 disrupting the proton wire. The structures of SE A227D at pH 4.5 and pH 8.0 are nearly identical showing a direct interaction with D147 which has replaced the water molecule in the proton wire. The β-sheet in SE A227D does not start until V150 causing H148 to reside on the external surface thus providing flexibility for D147 to interact with the chromophore. The other oxygen in the carboxylate side chain of D147 is exposed to solvent providing a pathway for external conditions to affect the chromophore. Despite SE A227D having a chromophore composed of T65, the proton wire connecting the chromophore to E222 is intact. Also note the position of T203 rotates in response to pH for GFP (S65T) while in SE A227D it does not. Oxygen molecules are in red. Nitrogen molecules are shown in darker blue. β-sheet is denoted as a broad ribbon in the protein backbone of the structure.

Two observations suggest that the pathway(s) conveying information to the chromophore of the FP domain are altered when the VSD responds to membrane potential transients. The first involves the positioning of the negative charge on the external surface of the FP domain. Moving the negative charge originally located at A227D along the exterior of the 11th β-sheet of the FP domain inverted the polarity of the voltage-dependent optical signal [9]. The position of that external negative charge determined whether the GEVI got dimmer or brighter upon depolarization of the plasma membrane. The VSD in the plasma membrane has been shown to dimerize [10] enabling intermolecular Förster Resonance Energy Transfer (FRET) between heterodimers of ArcLight-derived GEVIs having a FRET donor FP or a FRET acceptor FP [11]. Dimerization via the VSD could therefore bring neighboring FP domains into close proximity allowing the negative charge of the external A227D mutation in one protein to potentially commandeer the pH response of a neighboring chromophore.

The second observation suggests that the optical response of ArcLight involves changes to the internal environment of the FP domain. An ArcLight-derived GEVI with the internal E222H mutation to the FP domain exhibited a voltage-dependent increase in the 505 nm emission wavelength when excited at 390 nm that was not seen when the chromophore was excited at 470 nm [9]. Since the protonated chromophore excited at 390 nm requires a proton wire for deprotonation and subsequent emission at 505 nm, the voltage-dependent increase in emission upon 390 nm excitation revealed a temporal restoration of the proton wire enabling the long Stokes shift of the chromophore emission.

To enhance the modulation of the fluorescence, we have sought to investigate potential pathway(s) that enable external environmental conditions to influence the internal chromophore. ArcLight-derived GEVIs are ideal for this purpose as multiple (and repeatable) conformational changes can be induced with millisecond temporal precision via whole-cell voltage clamp of the plasma membrane. Exploring these pathways, we have found that internal amino acid sidechain orientation can be influenced by the immediate neighboring amino acids which due to the architecture of the β-sheet are located on the external surface of the protein. This interplay between surface chemistry and the internal environment of a fluorescent protein provide a novel approach for generating genetically encoded biosensors.

From these observations, we developed a new GEVI (Ulla) that gets substantially brighter upon depolarization of the plasma membrane thereby improving the contrast for detecting neuronal activity *in vivo*. Ulla clearly reported population activity in response to electrical stimulation in slice and odor stimulation in the olfactory bulb *in vivo*. The crystal structures of SE, SE A227D (ArcLight variant) and the FP domain of Ulla at different pH conditions also demonstrate potential flexibility of internal sidechains enabling threonine to be used as molecular switches to modulate hydrogen bonding inside a protein. Ulla should therefore be a useful tool for monitoring voltage transients in neuronal circuits and sets the stage for optimizing pathways in other FPs facilitating future biosensor development.

## Materials and Methods

### Plasmid design and construction

The Triple Mutant GEVI is described in Piao et al., 2015 [14]. The ArcLight construct is described in Jin et al., 2012 [8]. Primers were designed to introduce the point mutations to the FP as required. Conventional one-step and two-step PCR were used to generate the target inserts. The inserts were cloned into the vector using NEB restriction enzymes. The backbone vector used in this study is pcDNA3.1 (Invitrogen) with a CMV promoter.

### Cell culture and transfection

HEK 293 cells were cultured in Dulbecco’s Modified Eagle Medium (DMEM; Gibco, USA) supplemented with 10% Fetal Bovine Serum (FBS; Gibco, USA) in a 37 °C, 100% humidity and 5% CO2. For transfection, HEK 293 cells were suspended using 0.25% Trypsin-EDTA (Gibco, USA) then plated onto poly-L-lysine (Sigma-Aldrich, USA) coated #0 coverslips (0.08-0.13 mm thick and 10 mm diameter; Ted Pella, USA) at 70% confluency. Transient transfection was carried out with Lipofectamine 2000 (Invitrogen, USA) according to the manufacturer’s protocol.

### Electrophysiology

Coverslips with transiently transfected cells were inserted into a patch chamber (Warner instruments, USA) sealed with a #0 thickness cover glass for simultaneous voltage clamp and fluorescence imaging. The chamber was kept at 34 °C throughout the experiment and perfused with extracellular solution (150 mM NaCl, 4 mM KCl, 1 mM MgCl_2_, 2 mM CaCl_2_, 5 mM D-glucose and 5 mM HEPES, pH = 7.4). Filamented glass capillary tubes (1.5 mm/0.84 mm; World Precision Instruments, USA) were pulled by a micropipette puller (Sutter, USA) prior to each experiment to pipette resistances of 3–5 MΩ for HEK 293 cells. The pipettes were filled with intracellular solution (120 mM K-aspartate, 4 mM NaCl, 4 mM MgCl_2_, 1 mM CaCl_2_, 10 mM EGTA, 3 mM Na_2_ATP and 5 mM HEPES, pH = 7.2) and held by a pipette holder (HEKA, Germany) mounted on a micromanipulator (Scientifica, UK). Whole cell voltage clamp and current clamp recordings of transfected cells were conducted using a patch clamp amplifier (HEKA, Germany).

### Fluorescence microscopy of cultured cells

An inverted microscope (IX71; Olympus, Japan) equipped with a 60X oil-immersion lens, 1.35-numerical aperture (NA), was used for epifluorescence imaging. The light source was a 75 W Xenon arc lamp (Osram, Germany) placed in a lamp housing (Cairn, UK). GFP was imaged using a filter cube consisting of an excitation filter (FF02-472/30-25), a dichroic mirror (FF495-Di03) and an emission filter (FF01-496) for the 470 nm wavelength excitation all from Semrock (USA). Two cameras were mounted on the microscope through a dual port camera adapter (Olympus, Japan). A slow speed color charge-coupled-device (CCD) camera (Hitachi, Japan) was used to visualize cells during patch clamp experiments. Fluorescence changes of the voltage indicators were typically recorded at 1 kHz frame rate by a high-speed CCD camera (RedShirtImaging, USA). All the relevant optical devices were placed on a vibration isolation platform (Kinetic systems, USA) to avoid any vibrational noise during patch clamp fluorometry experiments.

### Optical data acquisition and analysis

Resulting images from patch clamp fluorometry were acquired and analyzed for initial parameters such as fluorescence change, ΔF = F_x_−F_0_ (F_x_ being the fluorescence during frame x, and F_o_ being the average fluorescence of the first five frames of the recording) or fractional fluorescence change values (ΔF/F = ((F_x_−F_0_)/F_0_) * 100) by NeuroPlex (RedShirtImaging, USA) and Microsoft Excel (Microsoft, USA). The acquired data from whole cell voltage clamp experiments of HEK 293 cells were averaged for 16 trials (technical replication) unless otherwise noted. The number of recorded cells in this work is biological replicates and the number of trials averaged for each HEK 293 cell should be interpreted as technical replication. Data were collected from recorded cells that did not lose their seals during the whole cell voltage clamp recording. ΔF/F values for the tested voltage pulses were plotted in OriginPro 2016 (OriginLab, USA). A ΔF/F trace versus time graph for each cell was also fitted for either double or single exponential decay functions in OriginPro 2016 as described previously (Piao et al., 2015).

### Protein expression and purification

The SE, SE A227D, and Ulla proteins were expressed by induction with 0.1 mM IPTG at 18 °C overnight. Cells were harvested by centrifugation and re-suspended in buffer containing 20 mM HEPES (pH 7.5), 150 mM NaCl, 2 mM β-mercaptoethanol, and were disrupted by sonication, and the crude lysate was centrifuged at 18,000 rpm (Hanil Supra 22K) for 40 min at 4 °C and the cell debris was discarded. The supernatant was loaded onto a nickel-chelated Hi-trap column (GE Healthcare) and eluted with a linear gradient of 25-500 mM imidazole in 20 mM HEPES (pH 7.5), 150 mM NaCl. The pooled fraction was further purified by gel filtration chromatography on HiLoad 26/60 Superdex-75 (GE Healthcare) pre-equilibrated with 20 mM HEPES (pH 7.5), 150 mM NaCl, and 2 mM DTT. The purity of the proteins at each stage was checked on 15 % SDS-PAGE. The purified SE, SE A227D, and Ulla proteins were concentrated to 35 ∼ 50 mg/ml using a VIVASPIN20 (Sartorius) concentrator and stored at −80 °C until use.

### Crystallization, Diffraction data collection and processing

To obtain pH-variation of the SE crystal, purified SE was equilibrated in 20 mM HEPES (pH 7.5), 150 mM NaCl, and 2 mM DTT at 4 °C for 2 hr prior to the crystallization attempts. The acid conditions of SE crystals were obtained by mixing an equal volume of the protein solution and a reservoir solution of 0.1 M sodium acetate (pH 4.5 ∼ 6.5) and 2.2 M Sodium Chloride. The neutral conditions of SE crystals were obtained by mixing an equal volume of the protein solution and a reservoir solution of 0.1 M HEPES (pH 6.5 ∼ 8.0), 2.5M sodium chloride, 12% PEG 1500 and 2.2 M sodium chloride. The basic conditions of SE crystals were obtained by mixing an equal volume of the protein solution and a reservoir solution of 0.1 M sodium Tris (pH 8.0 ∼ 9.5) and 0.8 M sodium dihydrogen phosphate. The acid conditions of SE A227D crystals were obtained by mixing an equal volume of the protein solution and a reservoir solution of 0.1M Citric acid (pH 4.5 ∼6.0) and 20%(w/v) PEG 6000. The neutral conditions of SE A227D crystals were obtained by mixing an equal volume of the protein solution and a reservoir solution of 0.1M Tris (pH 6.0 ∼ 8.0), 0.2 M lithium chloride and 20% PEG 6000. The basic conditions of SE A227D crystals were obtained by mixing an equal volume of the protein solution and a reservoir solution of 0.1M CHES (pH 8.0 ∼9.5), 0.2M sodium chloride and 1.26 M ammonium sulfate. The acid conditions of Ulla crystals were obtained by mixing an equal volume of the protein solution and a reservoir solution of 0.1 M acetate (pH 4.5 ∼ 6.5), 25% (w/v) PEG 1500 and 30% (w/v) MPD. The neutral conditions of Ulla crystals were obtained by mixing an equal volume of the protein solution and a reservoir solution of 0.1M Imidazole (pH 6.5 ∼ 8.0), 40% (w/v) Isopropanol and 15% (w/v) PEG 8000. The basic conditions of Ulla crystals were obtained by mixing an equal volume of the protein solution and a reservoir solution of 0.1M Tris (pH 8.0 ∼9.5), 0.1M Magnesium chloride and 25% (w/v) PEG3350. Suitable crystals for X-ray diffraction were grown within 5–7 days. They were cryo-protected using the reservoir solution supplemented with 20% (w/v) additional glycerol, ethylene glycol (w/v) and were flash-frozen in liquid nitrogen. Diffraction data were collected at −173 °C using the beamline BL-5C equipped with an ADSC Quantum 315r CCD detector of Pohang Light Source (Pohang, Korea). All data sets were processed and scaled using the program DENZO and SCALEPACK from the HKL2000 program suite [15].

### Structure Determination and Analysis

The electron density map was obtained by the molecular replacement method with the PHENIX[16] program using the crystal structure of GFP (PDB code: 5WWK) as a search model (ref: (2019) J Phys Chem B 123: 2316-2324). The model building and refinement were performed with Coot[17] and PHENIX[16], respectively. The final models were validated using PROCHECK [18]. The structure figure was visualized with PyMOL (http://pymol.org/, accessed on 3 May 2022). Supplementary table 2 summarizes statistics on the data collection and refinement.

### Viral injection and slice preparation

Virus injection and slice preparation were done according to an approved animal experiment protocol by the Institutional Animal Care and Use Committee at KIST (animal protocol 2019-099). 4-week-old C57BL/6N male mice were anesthetized by 1.5%-3% isoflurane. Adeno-associated virus (AAV) AAV2/1-hSyn-Ulla was then injected into the CA1 region of the hippocampus bilaterally as described previously (Lowery & Majewska, 2010). Recordings were performed at least 2 weeks after the injection.

300 mm thick acute coronal brain slices were prepared following the NMDG protective recovery method (Ting et al., 2018). Halothane (Sigma, USA) was used to anesthetize the mice before euthanasia. The isolated mouse brain was sliced in ice-cold high-sucrose artificial cerebrospinal fluid (ACSF) solution (75 mM Sucrose, 25 mM NaHCO_3_, 2.5 mM KCl, 0.5 mM CaCl_2_, 7 mM MgCl_2_, 1.25 mM NaH_2_PO_4_, 87 mM NaCl and 25 mM D-glucose) gassed with 95% O_2_/5% CO_2_ to give a pH of 7.4 using the VT-1200 Vibratome (Leica, Germany). Slices were then allowed to recover for at least an hour in a chamber filled with ACSF, incubated at 36 °C.**Slice imaging in dorsal hippocampal CA1 region.** Coronal slices of BL6 mouse brain injected with AAV2/1-hSyn-Ulla 2 weeks prior to the experiment were prepared following the NMDG protective recovery method, and the dorsal hippocampal CA1 region of the BL6 mouse expressing Ulla was imaged during a Schaffer collateral stimulation using bipolar electrodes (FHC, Bowdoin, ME). Brain slice recordings were carried out at 33 °C with the recording chamber continuously perfused with ACSF. The stimulus timing trigger and recording were done using Multiclamp 700B amplifier (Molecular devices, USA). Each Ulla recording was either acquired at 1 kHz as 80 pixel x 80 pixel or at 5 kHz as 26 pixel x 26 pixel fluorescence intensity time-lapse images using an upright epifluorescence microscope (Slicescope, Scientifica, East Sussex, UK) equipped with a High-speed CCD camera (Neuro CCD, RedShirtImaging, Decatur, GA), a GFP filter set (GFP-3035D-OMF, Semrock, Rochester, NY), a 10x water immersion objective lens (UMPlanFL N; NA=0.3, Olympus, Tokyo, Japan), and a 1.3 mW/cm^2^ 460 nm LED light source (UHP-Mic-LED-460, Prizmatix, Givat-Shmuel, Israel). The fluorescence intensity trace of Ulla was low pass filtered offline using a 50Hz 5th order Butterworth filter. The baseline prior to stimulation was used to apply an exponential decay curve. Traces represent a single recording or the average of 5, 10, or 20 recordings were averaged to derive the average Ulla response. The frame subtracted Ulla intensity images were pseudocolored by manually setting the colored intensity range from +0.3 %ΔF/F to −0.2 %ΔF/F and spatially low pass filtering with a 3 pixel x 3 pixel mean filter.

## In vivo imaging

### Surgical procedures

All experimental procedures were approved by the Florida State University ACUC. C57BL/6J adult male and female mice were anesthetized with ketamine/xylazine (90/10 mg/kg). Anesthetic depth was monitored during all surgical procedures via pedal reflex and a body temperature of 37.5 C was maintained via a heating pad placed underneath the animal. Mice were given a pre-operative dose of carprofen, dexamethasone, atropine and bupivacaine, and Lacrilube was applied to their eyes. For virus injections, an incision was made in the skin above the olfactory bulb, a small craniotomy was made and 500 nl of AAV2/1-hSyn-Ulla was injected into the olfactory bulb over a period of 15 minutes (Nanoject III, Drummond Scientific). Following completion of the injection, the incision was sutured and mice were administered post operative doses of rimadyl for 3 days. A minimum of 2 weeks were allowed for Ulla expression before performing imaging experiments in which similar surgical procedures were used to attach a head-post to the skull using metabond (Parkell, Edgewood, NY). The skull above one or both olfactory bulbs was thinned and covered with cyanoacrylate to improve optical clarity or was removed and sealed with a glass coverslip.

### Imaging

Imaging was carried out using a custom epifluorescence microscope built from Thorlabs (Newton, NJ) components with a 35 mm or 50 mm CCTV lens, or a 10x 0.5 N.A. microscope objective. Illumination was provided by a Prizmatix LED (UHP-T-LED-455) filtered with a 488/10 nm (Semrock FF01-488/10). A dichroic mirror (59009bs, Chroma) directed the excitation light towards the preparation, and transmitted the fluorescence emission through a band-pass filter (Chroma 59009m), before being recorded using a DaVinci 1k camera (RedShirtImaging, Decatur, GA). Imaging was carried out between 40-200 Hz at a spatial resolution of 256×256 pixels. 2-photon imaging was carried out on a Sutter Moveable Objective Microscope (MOM, Sutter Instruments) equipped with an 8 kHz scanner (Cambridge), GaAsP photomultiplier tubes (Hamamatsu), and a 2-photon optimized objective (10x 0.5 NA). A Sparks Lasers ALCOR-920 provided 2-photon laser excitation which has an internal AOM to control its power. This system acquires images at 30.9 Hz at a spatial resolution of 512×512 pixels. During imaging experiments, mice were placed underneath the microscope objective during imaging with an odor delivery port placed in front of their nose.

### Olfactometer

The odor methyl valerate (acquired from Sigma-Aldrich) was presented to the animal using a custom flow-dilution olfactometer at concentrations between 1-10 % of saturated vapor. In some preparations respiration was measured using a pressure sensor attached to the olfactometer odor delivery port.

## Data analysis

Odorant-evoked signals were collected in consecutive trials separated by a minimum of 60 seconds. Ulla signals representing depolarization are shown as upward signals. Data from each pixel were low-pass Gaussian filtered between 2-4 Hz and had an exponential drift subtracted that was calculated based on the signal prior to the stimulus onset (exponential subtraction function in NeuroPlex). The signal from each pixel was divided by its resting fluorescence measured at the beginning of each trial. The fluorescence brightness analysis was measured by choosing a region of interest that captured the entire surface of the injected and uninjected hemibulbs. The mean fluorescence in the initial 2 seconds of the imaging trial was measured in both hemibulbs in 4 trials and averaged together. The results were normalized to the fluorescence in the uninjected hemibulb and averaged across all four preparations. The ΔF/F analysis was carried out by subtracting a 1-s window around the peak of the odor response from the 2-s immediately preceding the odor stimulus. This measurement was carried out in 4-5 trials per preparation before being averaged together. The 2-photon imaging data were analyzed in spatially and temporally averaged files in which the pixel resolution was reduced to 256×256 and in which every 4 frames were averaged together to improve the signal-to-noise ratio using MATLAB (Mathworks) and MView (Sutter instruments). The 2-photon fluorescence traces were generated from an averaged file in which all single trials were aligned to the first inhalation following the odor command trigger. The *in vivo* 2-photon mean fluorescence is from the original 512×512 resolution data. All optical traces were generated from the spatially and temporally averaged files, although similar signals were observed in the raw data files. Data analysis was performed using Turbo-SM analysis software (SciMeasure, Decatur GA), and images were arranged using Adobe Illustrator (Adobe).

## Results

To couple fluorescence changes with membrane potential transients, ArcLight-derived GEVIs fuse SE A227D to a VSD. Figure 1A shows the Alphafold prediction of the GEVI Triple Mutant (TM, an ArcLight-derivative with faster kinetics [14]). When the voltage across the plasma membrane changes, the VSD will alter its conformation causing the FP to also move. How the FP movement causes a change in fluorescence is not clear. One hypothesis suggests the possibility of the FP domain being distorted as it moves against the plasma membrane [19]. This is unlikely since other FPs fused to the VSD do not yield large, voltage-dependent optical signals [20]. Indeed, SE also requires an additional external mutation (A227D) to exhibit a large voltage signal. The original SE yields voltage-dependent signals of ∼1% ΔF/F/100 mV while SE A227D yields ∼40% ΔF/F/100 mV [8, 21].

An alternate hypothesis involves the interaction between the external sidechains of one FP domain with the external A227D sidechain of a neighboring GEVI causing the optical response to voltage transients [22]. The possibility of nearby FP domains interacting was demonstrated by a voltage-dependent intermolecular Fluorescence Resonance Energy Transfer (FRET) signal when separate GEVIs having a FRET donor or acceptor FP are co-transfected into the same cell [20]. This interaction between FP domains is possible via the dimerization of the voltage-sensing phosphatase gene family [10].

### A titratable sidechain in the β-can connects the external environment to the chromophore

The interaction between neighboring FP domains of the GEVI suggests that the A227D mutation creates an external negative field that interacts with a receptor region on a separate, neighboring FP domain. When the VSD moves, the relationship between neighboring FP domains is altered. In essence, controlling membrane potential enables the rapid movement of a negative field near a neighboring FP domain. We therefore crystallized SE A227D and SE in different pH conditions in an attempt to reveal changes in the FP that could mediate fluorescence change (Figures 1B and 1C).

SE is an extremely pH-sensitive FP [23] which is likely due to the orientation of an aspartic acid sidechain (D147) in the can structure spanning from the external surface of the FP to the internal chromophore. One oxygen of its carboxylate sidechain is exposed to the external surface while the other oxygen forms a hydrogen bond to the chromophore (Figure 1B). A similar structure was recently reported for the FP, Lime, a derivative of SE that improves upon the pH sensitivity of the FP [24]. In contrast, the EGFP structure shows the hydroxyl group of S147 exposed to the exterior of the protein [12].

Closer inspection near the chromophore of the FP structures reveals a similar hydrogen bond network to that of the original GFP (avGFP, Figure 1C) with subtle differences in the secondary structure of the 7^th^ β-sheet as well as the orientation of the T203. The 7^th^ β-sheet of SE A227D does not start until V150 while in avGFP the β-sheet starts at H148. As a result, the sidechain of H148 now resides on the external surface while D147 has sufficient space to interact with the chromophore.

Remarkably, despite the presence of T65 in the SE A227D chromophore, the proton wire connecting the chromophore to E222 is intact. In GFP (T65S), the E222 sidechain interacts with T65 and no longer interacts with S205 causing a break in the proton wire (Figure 1C). However, in SE A227D, E222 does not interact with T65 and forms a hydrogen bond with S205. D147 replaces the water molecule in the avGFP proton wire providing a pathway for external pH to affect the pKa of the chromophore as well as alternate routes of proton migration inside the protein.

Of further note is the orientation of the T203 sidechain. When the chromophore is protonated, T203 adopts a configuration where the hydroxyl group is oriented away from the chromophore (Figure 1C) [6, 13]. For GFP (S65T), at alkaline pH 8.0, T203 rotates the sidechain allowing the hydroxyl group to interact with the anionic chromophore [7, 13, 25]. For SE A227D, T203 adopts an orientation similar to avGFP and GFP (S65T) in low pH but does not rotate in response to alkaline pH conditions. This rigid orientation of T203 is true for the original SE structure over a broad pH range as well (see Supplemental figure S1). D147 therefore appears to be the primary cause of pH response. Indeed, when S147D mutation was introduced into EGFP, the pKa was shifted to 7.2 [26].

### D147 is not directly involved in the voltage-dependent optical signal of the GEVI

With its sidechain spanning the external surface of the protein to the internal chromophore, D147 was a likely candidate for interacting with the A227D sidechain of a neighboring FP domain. However, for the voltage-dependent optical response, the D147A mutation had very little effect compared to the original TM response (Figure 2B). Other mutations to D147, however, had a distinct effect on the voltage-dependent signal. For instance, D147K resulted in a multicomponent optical response with an initial decrease in fluorescence followed by a slower increase during the voltage step. Upon the return to the holding potential, the fluorescence overshoots the initial fluorescence before returning to baseline. The same pattern was seen for the D147N mutant but not the D147Q which resembled the original D147 response.

**Figure 2.**
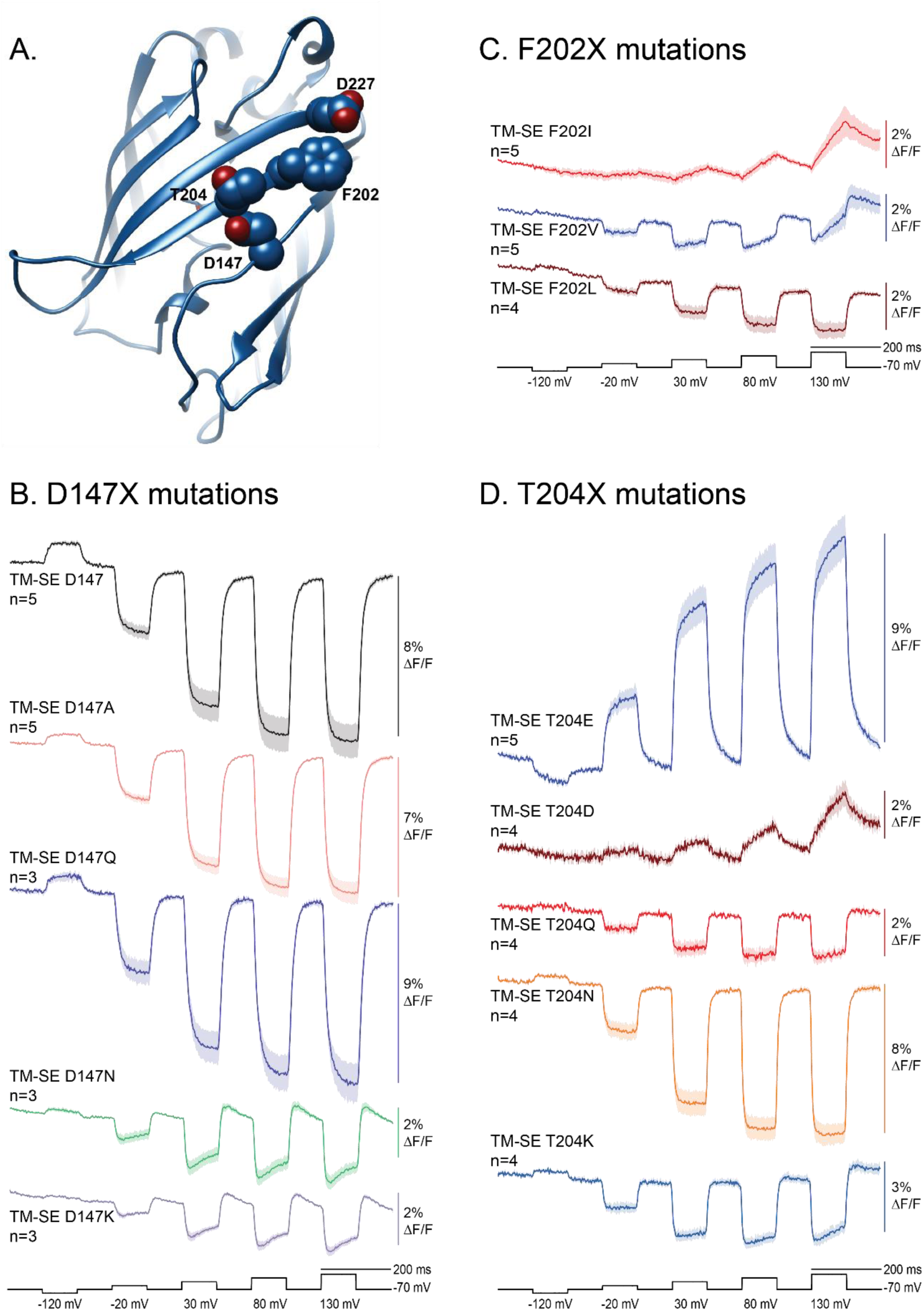
Altering the voltage-dependent optical transitions via mutations to the externally exposed sidechains of the FP domain. A. Position of the four amino acids implicated in the large voltage-dependent optical signal of ArcLight-derived GEVIs. The molecular positions of the sidechains for D147, F202, T204, and D227 are shown in relation to the β-sheet backbone as determined by the crystal structure of SE-A227D pH 8.0 (Figure 1B). **B. Optical traces of HEK cells expressing the original GEVI, TM-SE D147, or GEVIs mutated at the D147 position in the FP domain.** Under whole cell voltage clamp conditions, cells were held at −70 mV holding potential and subjected to voltage steps as indicated in the black trace below. N represents number of cells recorded. Each cell was subjected to 16 repetitions of the voltage step protocol. The solid line is the average of the mean. Shaded area represents the standard error. Acquisition frame rate was one kHz. An offline butterfield filter 100 Hz was applied to all traces unless otherwise stated. **C. Effect of nonpolar, nonbulky sidechains at the 202 position.** The F202 position was changed to either isoleucine (red trace), valine (blue trace), or leucine (dark red trace). **D. Amino acid sidechain charge and/or orientation affect the voltage-induced fluorescence transitions.** A negative charge at position 204 inverts the voltage-dependent signal with glutamic acid having a larger dynamic range than aspartic acid. The polar amino acids glutamine and asparagine also exhibit differences in the dynamic range with T204Q resembling the T204K mutant, while T204N resembles the response of the original GEVI (shown in B.).

While these results ruled out the carboxylate sidechain of D147 as the potential receptive field for the interaction of the A227D charge from a neighboring FP domain, they also demonstrated the flexibility of 147 position. There are too many steric conflicts for lysine to adopt the same structure as the original D147 suggesting alternate conformations of the 7^th^ β-strand depending on the D147X mutation. In addition, the distinct optical responses of D147N versus D147Q suggested a role for the orientation of the sidechain as asparagine and glutamine have the same functional carboxyamide sidechain differing in length by a single carbonyl. The fact that the amide of the lysine from the asparagine at position 147 generated a multicomponent voltage response but the amide for glutamine at 147 had no effect suggested that the orientation of the functional units at position 147 influenced the voltage-dependent signal.

### Hydrophobicity plays a role in the voltage-dependent optical response

D147, F202, and T204 have been shown to be required for the large voltage-dependent optical signal of ArcLight in addition to the serendipitous A227D mutation [8]. In GFP, the polar amino acid, serine, resides at position 202 suggesting that, for the optical transitions, external nonpolar residues also play a role in directing information from the surface of the FP to the internal chromophore. We therefore introduced nonpolar, non-aromatic amino acids to the F202 position of the FP domain. Remarkably, the hydrophobic architecture of the sidechain at the 202 position had a dramatic effect on the voltage-dependent optical signal (Figure 2C).

A secondary component to the voltage-dependent optical signal was seen for two of the hydrophobic changes at the F202 position. The F202V mutant had a reduced response below 1% ΔF/F for the 50 mV depolarization step (Figure 2C). Stronger depolarization of the plasma membrane caused the introduction of a secondary optical response, the fluorescence shows a fast decrease in fluorescence followed by a slow increase much like the response seen for D147N and D147K constructs (compare Figures 2B and 2C). However, unlike D147N and D147K, the secondary response for F202V becomes more pronounced as the depolarization step is increased. The F202I construct also shows an increase in the secondary response that becomes more pronounced as the depolarization step of the plasma membrane is increased, but the initial decrease in fluorescence for F202I is nearly undetectable. This is rather surprising given that the F202L mutant only exhibited the initial decrease. F202L favored the initial decrease while F202I favored the slower increase.

## Sidechain charge at the external 204 position in the FP domain inverted the voltage-dependent optical signal

The effect of different hydrophobic amino acids at position 202 has interesting implications. There are a significant number of hydrogen bonds along the surface of the β-barrel of GFP providing multiple avenues of potential charge displacement routes along the surface of the protein [3, 27]. The phenylalanine at position 202 could shield and/or isolate the polar sidechain of the adjacent external T204 position (Figure 2A). Since T204 was also shown to be required for the large voltage-dependent optical signal of ArcLight [8], we introduced different polar and charged amino acids at position 204.

Mutations to the T204 position revealed that sidechain charge as well as orientation affected the voltage-dependent optical signal (Figure 2D). When a negative charge was introduced at the 204 position, the voltage-dependent optical signal inverted slightly for the T204D mutation (Figure 2D) while a more robust inversion of the optical signal was seen for the T204E mutant (+9% ΔF/F/200 mV). The T204R yielded only a 2% maximal dimming of the fluorescence upon a 200 mV depolarization of the plasma membrane with a multicomponent response similar to some of the D147X and F202X mutations, a fast decrease in fluorescence followed by a slower increase. Again the response of the asparagine replacement differed from the glutamine response. This time the T204Q construct responded to voltage transients with a multicomponent optical signal while the T204N resembled the original TM GEVI (compare Figures 2B and 2D).

## The mechanism mediating the voltage-dependent optical response involved an internal conformational change near the chromophore in the FP domain

The effects on the voltage-dependent optical signal for the F202 and T204 mutations (Figures 2C and D) implicated T203 as a potential conduit relating external conditions to the chromophore. It is well established that the 203 position can affect the ground state fluorescence of the protein with non-polar replacements at 203 favoring the protonated chromophore and T203Y red-shifting the emission spectrum of the fluorescence for YFP [25, 28, 29]. We therefore initially examined T203X mutations in the presence of T204E to ask if T204E would always cause the fluorescence to increase upon depolarization of the plasma membrane regardless of the amino acid at 203.

Fluorescence was observed in the double mutants T203E/T204E, T203Q/T204E, T203N/T204E, and the T203H/T204E (Figure 3B). T203D/T204E, T203R/T204E, and the T203K/T204E double mutants did not fluoresce (see supplemental figure S2). The T203E/T204E and the T203Q/T204E constructs behaved very much like the T204E single mutation by exhibiting an increase in fluorescence upon depolarization of the plasma membrane (compare Figures 2D and 3B). The T203N/T204E also got brighter during the depolarization steps but at a reduced level. Interestingly, the T203H/T204E double mutant dimmed upon depolarization of the plasma membrane. The 203 position could counteract the influence of T204E requiring the need to explore the effects of T203X mutations.

**Figure 3.**
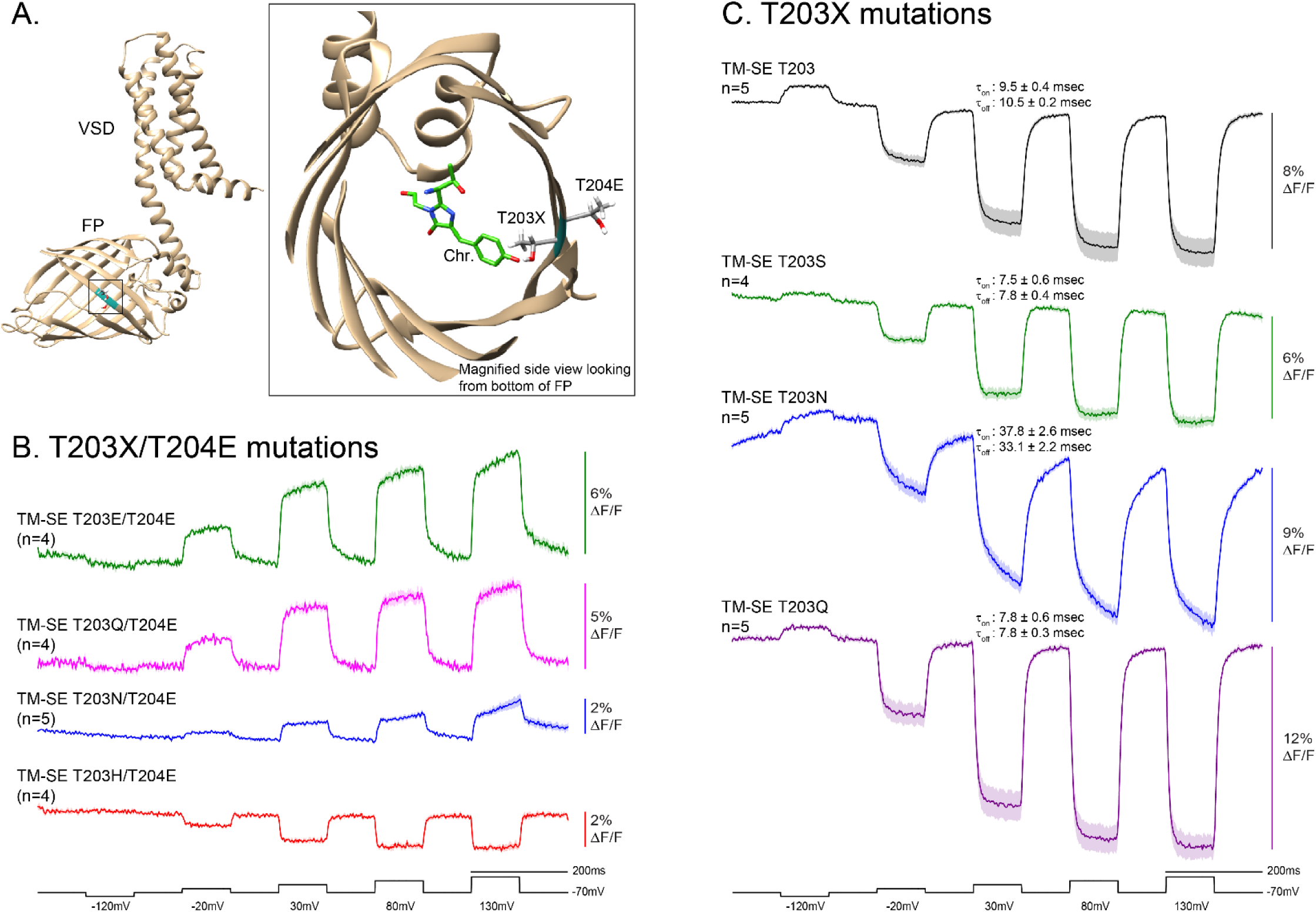
Interplay between the internal 203 position and the external 204 position in the FP domain of the GEVI. A. Schematic of TM-SE GEVI predicted by Alphafold. Schematic is the Alphafold prediction of the TM-SE GEVI with the voltage sensing domain (VSD) residing in the plasma membrane and the fluorescent protein domain (FP) in the cytoplasm. The random coil amino terminus has been removed for clarity. The positions colored aqua in the FP domain highlight the T203 and T204 amino acids. The side view looking from the bottom of the FP domain shows the internal orientation of the sidechain for T203 and the external orientation for the T204 sidechain. The chromophore (Green) is positioned based on the crystal structure of GFP as in figure 1. **B. Double mutations to the T203 position with T204E.** HEK cells expressing a T203X mutation in the presence of T204E (double mutants) were subjected to whole-cell voltage clamp and imaged at 1Khz. **C. Single mutations to the internal T203 site affected the speed of the voltage-dependent optical signal.** HEK cells expressing only T203X mutations (T203X/T204) were subjected to whole-cell voltage clamp and imaged at 1Khz. The top trace is the original TM GEVI from figure 2 shown for comparison. In the absence of the T204E mutation, T203N and T203Q both get dimmer during depolarization steps but at significantly different rates.

In the absence of the T204E mutation, all of the GEVIs with T203X single mutations got dimmer in response to depolarization of the plasma membrane (Figure 3C). There were three exceptions. The T203D, T203E, and the T203R single mutants did not fluoresce (Supplemental figure S2). The lack of fluorescence for T203E is noteworthy since the T203E/T204E mutant did fluoresce suggesting an interplay between the external 204 position and the internal 203 position. In addition, the T203K single mutant fluoresced while the T203K/T204E did not (Supplemental figure S2). The most striking observation, however, was that a single mutation to an internal side chain in the FP domain affected the speed of the voltage-dependent optical response of the GEVI. The signal onset for the original TM GEVI was best fit by a single exponential decay with a τ_on_ of 9.5 ± 0.5 msec. The return to baseline for the TM was also best fit by a single exponential decay (τ_off_ of 10.5 ± 0.2 msec, Supplemental table 1). TM-SE T203Q has a slightly faster onset than the original construct (τ_on_ ∼ 7.8 ± 0.6 msec). However, the TM-SE T203N mutant had a slowed optical response with the onset of the optical signal best fit by a double exponential decay. The fast τ_on_ for T203N (9.1 ± 0.7 msec) was similar to the original TM GEVI (9.5 ± 0.5 msec) but only accounted for 56.6 ± 0.1 % of the total signal. The slow τ_on_ T203N was 75.7 ± 5.8 msec. T203Q has a faster return to baseline with a minor secondary component with a τ_off_ best fit by a double exponential decay (fast τ_off_ ∼7.2 ± 0.2 msec comprising 94.3 ± 0.1 % of the total signal). T203N was again slower in returning to baseline with a slow τ_off_ of 69.7 ± 6.2 comprising 38.0 ± 0.1 % of the total signal.

## Chromophore flexibility can be offset by mutations to the surface chemistry of the FP domain: demonstration of a threonine switch

The altered kinetics by tens of milliseconds when only an internal amino acid was mutated suggested a conformational change near the chromophore occurring during the voltage-dependent optical signal. Given that the structures in Figure 1 show T203 is capable of adopting different conformations, we wondered if the rotation of T203 would alter the flexibility of the chromophore as well. The crystal structure data for SE A227D and SE revealed no obvious conformational changes in response to pH, but we have already shown that the pH response is distinct from the voltage-dependent optical signal. In addition, transiently affecting the flexibility of the chromophore would only be observed if intermediate conformations were stable enough to crystalize.

We therefore changed the question to what would happen to the voltage-dependent optical signal if we altered the flexibility of the chromophore. To do this, we compared the structure around the chromophore of SE A227D to a pH-sensitive protein possessing a chromophore that does exhibit different conformations in different pH conditions (Figure 4A). The red-shifted pH-sensitive FP, Katushka, has structure data at pH 5.5 (dim state) and pH 8.0 (bright state) [30]. An overlay of the structures for Katushka reveals that the chromophore is in the cis-configuration in the bright state, while the lower pH structure shows an altered configuration of the chromophore potentially interacting with the β-can at position S158. Comparing the structures of SE A227D with Katushka, however, revealed that the homologous S158 position in Katushka is phenylalanine (F165) which would sterically hinder the chromophore of SE from adopting the alternate configuration (Figure 4A top right). We therefore mutated the F165 position of SE A227D to serine, threonine, and tyrosine to explore the effects a potentially more flexible chromophore would have on the voltage-dependent optical signal.

**Figure 4.**
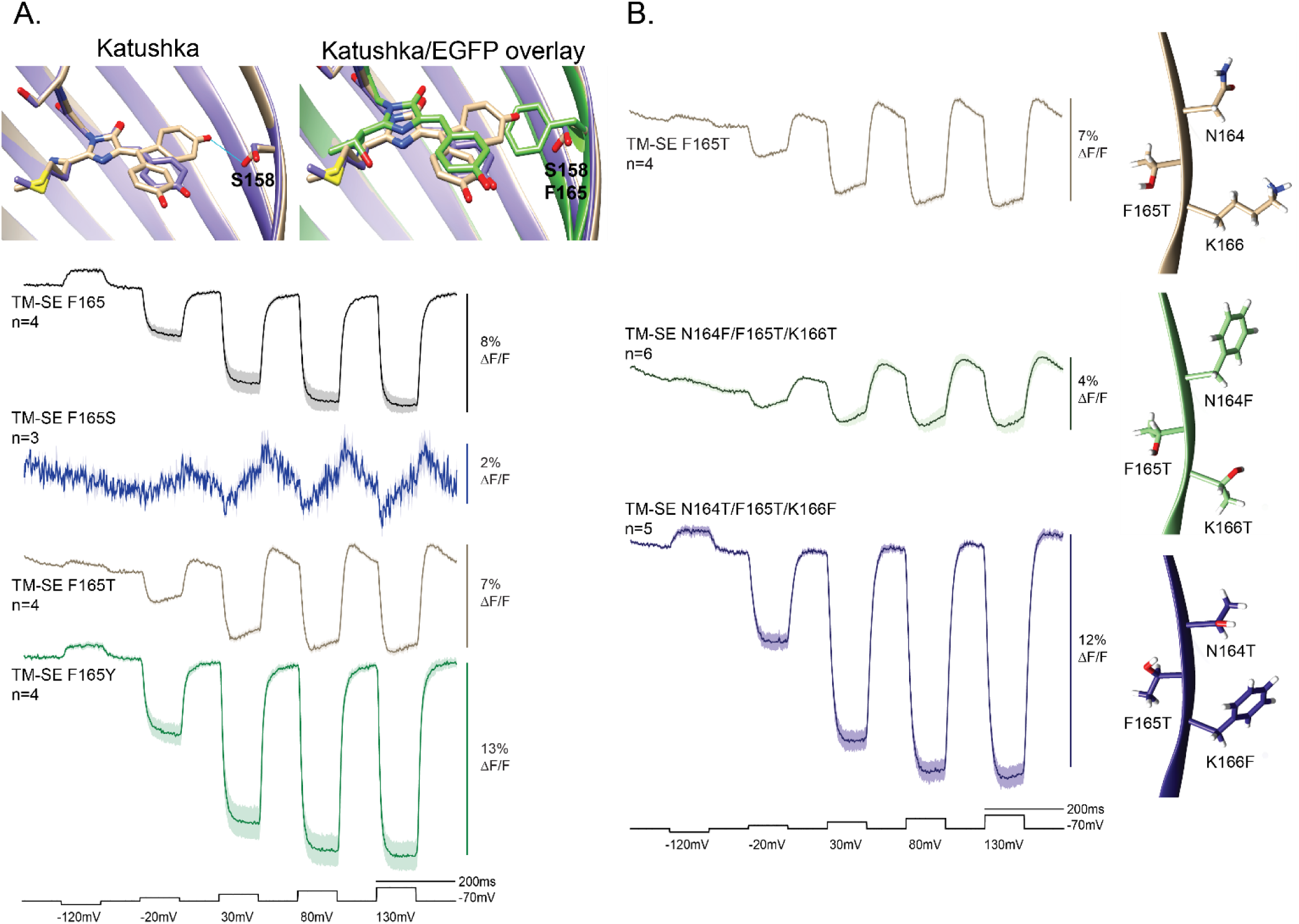
Compensating chromophore flexibility with mutations to the surface chemistry of the FP domain. A. Steric hindrance at the 165 position in the FP domain inhibits a multicomponent voltage-dependent response. Top left structure is an overlay of the pH-sensitive, red fluorescent protein, Katushka at pH 8 (bright state - purple structure) and pH 5.5 (dim state – beige structure) [30]. The dim state reveals a conformational change to the chromophore creating a potential hydrogen bond (blue line) with the hydroxyl group from serine 158. The top right structure is an overlay of the Katushka structures with GFP. The phenylalanine at the homologous 165 position of GFP would sterically inhibit a similar conformational change in SE A227D. Traces below are from HEK cells expressing the F165 mutant constructs and subjected to whole-cell voltage clamp. For comparison, the original TM (F165) is shown in black reproduced from figure 1. The voltage dependent signal from the F165S mutant is shown in blue. The F165T response is shown in brown, and the F165Y response is shown in green. **B. Asymmetrical polarity on the surface of the FP affects the fluorescence transition during depolarization of the plasma membrane.** The multicomponent optical response for the F165T mutant is the beige trace. The structure on the right is the Alphafold prediction. Mutating the external flanking amino acids (N164F/K166T) results in a similar multicomponent response with a lower dynamic range (green trace). Inverting the external polar asymmetry (N164T/K166F) increases the dynamic range of the optical response and nearly eliminates the secondary component during the transition in fluorescence. The Alphafold predictions suggest an alternate conformation for the threonine at the 165 position depending on the amino acids at 164 and 166.

Replacement of the bulky phenylalanine sidechain of SE A227D in the FP domain of the GEVI with serine produced a construct with dim fluorescence yet yielded a multicomponent optical signal similar to those seen in some of the D147X, F202X, and T204E mutations with a rapid decrease followed by a slower increase in fluorescence (Figure 4A). The F165T version is brighter than the F165S mutant yet exhibits the same multicomponent voltage-dependent optical response. The F165Y mutant is also bright but does not show the multicomponent response of the F165S or F165T mutants indicating that a bulky sidechain at 165 be it F165 or F165Y inhibits the multicomponent voltage-dependent transition. The phenotype of the bulky amino acids at the 165 position and the altered response for F165S and F165T suggest that the secondary component may be due to a more flexible chromophore.

Having a novel internal mutation (F165T) with a multicomponent voltage-dependent signal provided an opportunity to address how the external chemistry flanking the 165 position could alter the transition of the fluorescence in response to membrane potential transients. Given the potential role for hydrophobicity at the external F202 position and the effect of polarity at the external T204 position in the FP domain (Figure 2), we constructed two new GEVIs consisting of either the N164F/F165T/K166T (F_outside_/T_inside_/T_outside_) or the N164T/F165T/K166F (T_outside_/T_inside_/F_outside_) triple mutations to the FP domain (Figure 4B). The N164F/F165T/K166T triple mutant resembled the optical pattern of the F165T single mutation with a smaller change in fluorescence and a slower secondary component. Inverting the polarity across the 165 position (N164T/F165T/K166F) significantly altered the voltage-dependent signal nearly eliminating the secondary component (a slight secondary component can be seen in the 150 mV and 200 mV depolarizations steps) and doubling the size of the optical response. The external chemistry flanking the 165T determined the pattern of the fluorescence transition in response to voltage. The direction of the asymmetrical polarity at positions 164 and 166 determined if 165T behaved like 165S or the bulky 165F and 165Y; a functional demonstration of threonine behaving as a molecular switch.

The Alphafold predictions revealed variable orientations for the F165T side chain depending on the surface chemistry at the 164 and 166 positions (Figure 4B). In the F165T mutant the hydroxyl group was oriented towards the lysine at the 166 position. Introducing asymmetric polarity to the two flanking amino acid positions (N164F/F165T/ K166T) exacerbated the multicomponent pattern and showed a similar orientation for the internal threonine at the 165 position. However, Alphafold predicts that reversing the polarity of the surface chemistry flanking the 165 position (N164T/F165T/K166F) orients the hydroxyl group of 165T towards the 166 position. This conformation of 165T places the methyl group towards the 164 position potentially restoring the steric hindrance of the original F165 to some degree and thereby inhibiting/reducing the multicomponent voltage-dependent optical signal of the F165T mutant.

## Repositioning the external negative charge along the 11^th^ β-sheet for the T204E mutant yields a novel GEVI, Ulla (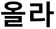) with an inverted voltage signal and an altered pH sensitivity

The surface chemistry of the FP domain influencing the orientation of internal amino acids near the chromophore has an interesting consequence; asymmetrical polarity on the surface of the FP domain creates external ‘hotspots’ capable of responding to changes in environmental conditions, i.e., external electric fields. We therefore explored the possibility A227D is not the ideal position for the negative charge in the presence of the T204E mutation. Previous attempts to relocate the negative charge on the exterior of the FP domain from the A227D position failed to improve the voltage-dependent optical signal for ArcLight-derivatives [9]. For the T204E mutant, however, relocating the external D227 negative charge more than doubled the dynamic signal. The negative charge at position D227 was reverted to alanine, and an aspartic acid scan of the remaining external residues along the 11^th^ β-sheet was performed (Figure 5A). The best voltage-dependent optical signal was the T204E/F223D yielding a fluorescence increase greater than 25% ΔF/F upon a 100 mV depolarization of the plasma membrane. This new GEVI was named Ulla (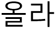-Korean meaning to ascend).

**Figure 5.**
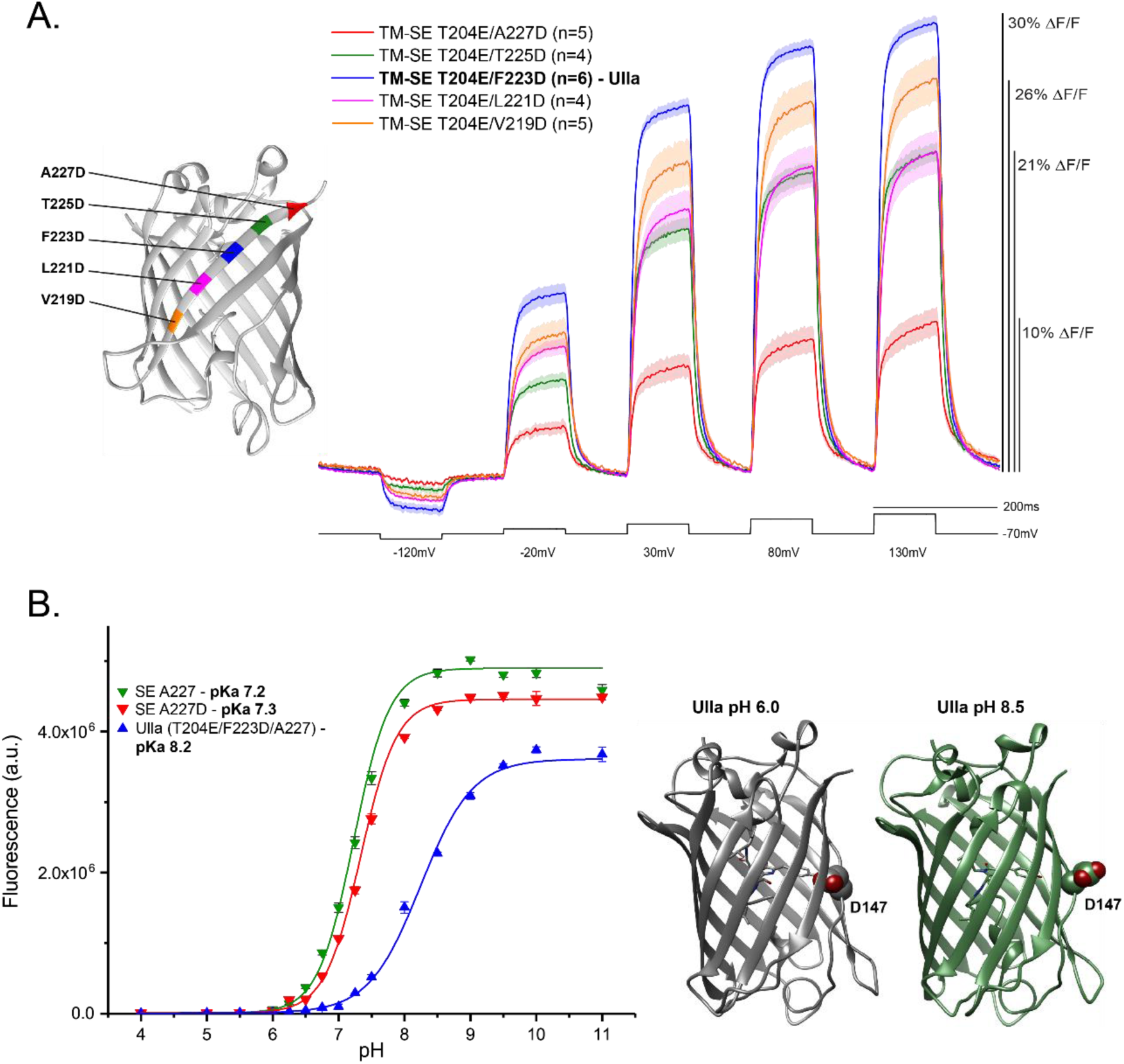
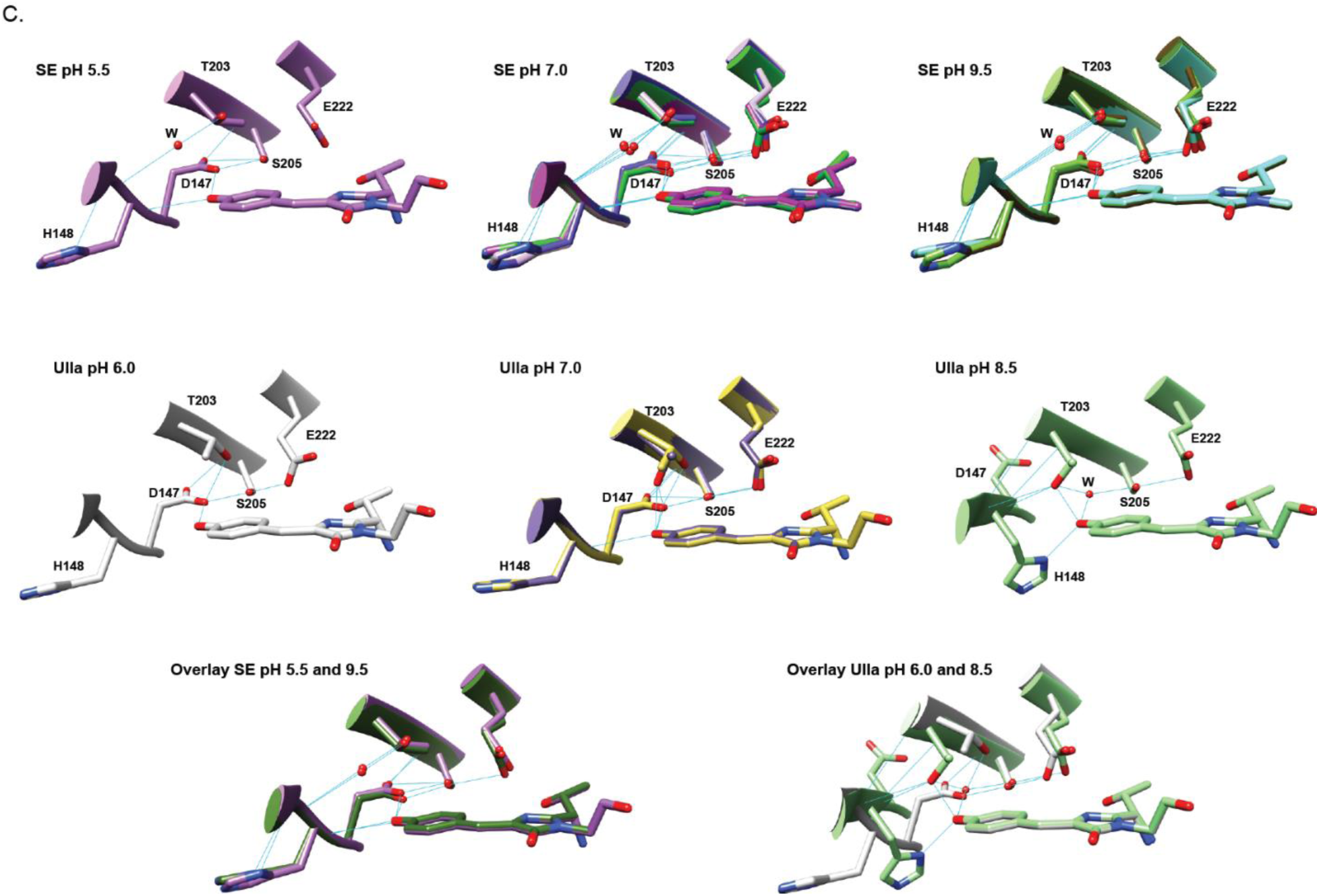
Influencing the orientation of T203 by mutations to the surface chemistry of the FP domain creates a novel GEVI with a shifted pH sensitivity. A. Aspartic acid scan on the external amino acid of the 11^th^ β-sheet in the presence of T204E. The 227 position in the FP domain was reverted back to alanine. An aspartic acid residue was then introduce at either position 225 (T225D – green trace), 223 (F223D – blue trace), 221 (L221D – pink trace), or 219 (V219D – orange trace). **B. The FP domain of Ulla (T204E/F223D) has a shifted pKa.** The FP domains of SE, SE A227D, and Ulla were purified from a bacterial expression system. The fluorescence of all three increases as the pH is raised with Ulla having a pKa outside the physiological range (8.2). **C. The external mutations in Ulla (T204E/F223D) alter the orientation and flexibility of the internal T203 sidechain near the chromophore compared to the parent SE structure.**The FP domain of Ulla and the original SE lacking the A227D mutation of ArcLight were also crystallized in different pH conditions. The structure of below the pKa (pH 6.0) resembles the original SE protein and the SE A227D with the D247 sidechain both exposed to the external surface as well as interacting with the chromophore. Above the pKa (pH 8.5) the 7^th^ β-sheet exhibits a conformational change in 3 of the 4 structural units. The fourth unit has the D147 sidechain still interacting with the chromophore (see PDB 8JLU). The structures in the top row are for the parent SE protein at pH 5.5, 7.0, and 9.5 respectively. Hydrogen bonds are depicted by blue lines. Oxygens are shown in red. Nitrogens are shown in blue. SE crystallized as four unit cells for each pH condition. Only one structure is shown for the SE pH 6.5 condition while all four structures are overlaid in the SE pH 7.0 and SE pH 9.5 conditions. For Ulla, the pH 6.0 condition yielded a single structural unit with T203 in an alternate orientation compared to the SE structures. The two structural units of Ulla at pH 7.0 are overlaid revealing a more flexible T203 sidechain. At pH 8.5, four structural units were solved for Ulla, three of which show an extension of the 7^th^ β-sheet resulting in D147 being solely on the external surface of the FP domain. A water molecule (W) has replaced the oxygen of the D147 sidechain that interacted with the chromophore at lower pH conditions. H148 has also rotated from the external surface and now interacts with the chromophore as well.

The altered surface chemistry of the FP domain for Ulla has the added benefit of shifting the pH sensitivity of the GEVI. The Ulla FP domain, the original SE, and the SE A227D FP domain of ArcLight were expressed in bacteria and purified to compare their pH sensitivities (Figure 5B). All three FPs increased their fluorescence as the pH was raised. SE and SE A227D had very similar responses (pKa of 7.2 for SE and 7.3 for SE A227D). However, the pKa for Ulla was shifted to a more alkaline pH of 8.2. Ulla differs from SE by only two external mutations, T204E/F223D, yet the pKa was shifted a full pH unit. Ulla is still pH sensitive, but in the physiological range of the cell that sensitivity is much reduced.

## Crystal structures of the Ulla FP domain revealed a pH-induced flexibility in the 7^th^ β-sheet as well as an internal threonine switch at position 203

Control of the threonine switch introduced to the 165 position (Figure 4) was inspired by the F202 and T204 mutations found in the original SE protein (Figure 1). With Ulla having a mutation to the 204 position and an altered pKa, the crystal structures of the FP domain of Ulla (SE T204E/F223D) in different pH conditions were solved (Figure 5B) and compared to the structures of the original SE protein (Figure 5C).

Ulla, shows two important conformational changes in response to pH. The first is the exclusion of D147 from the interior of the protein when the pH exceeds the pKa. At pH 6.5, D147 interacts with the chromophore in a manner similar to SE A227D and SE (Figure 5B). However, at pH 8.5, there is an extension of the 7^th^ β-sheet resulting in the carboxylate of D147 on the external surface of the β-can structure. In this configuration, a water molecule occupies the position vacated by the carboxylate sidechain of D147, maintaining the hydrogen bond pattern with the chromophore. It should be noted that one of the four structural units in the Ulla pH 8.5 does not extend the7th β-sheet resulting in D147 still interacting with the chromophore which may be a consequence of being near the pKa of the protein (See PDB 8JLU).

Another conformational change observed for Ulla is the orientation of the T203 sidechain providing a second demonstration of a threonine switch. For the original SE in pH conditions below, near, and above the pKa, the T203 hydroxyl group is oriented towards a water molecule bridging the gap between the backbone of the 7^th^ and 10^th^ β-strands likely stabilizing the can structure (Figure 5C). Indeed, the positioning of the chromophore, and the sidechains of D147, T203, and S205 are virtually invariant across a broad pH range. However, for Ulla, the orientation of T203 is much more flexible. When the pH is below the pKa for Ulla, T203 is a different orientation than that of SE enabling a hydrogen bond with the internal oxygen of D147 (Figure 5C). The increased flexibility of the T203 for Ulla can be seen as the pH is raised. At pH 7.0, Ulla exhibits 2 distinct orientations for the T203 sidechain and multiple orientations at pH 8.5 allowing for direct interaction with the chromophore. Recall that Ulla differs from SE at the 204 and 223 positions both of which reside on the external surface of the β-can structure.

## Population signals from the novel GEVI, Ulla

A GEVI that gets 25% brighter upon depolarization of the plasma membrane offers an exciting new tool for discerning the activity of neuronal circuits. Voltage imaging poses many challenges, not the least of which is having a functional probe that is restricted to the plasma membrane [31]. With the fluorescence distributed along the surface of neurons, the interspersion of neuronal processes generates a significant background fluorescence that can make it difficult to observe action potentials let alone synaptic inputs. Having a GEVI that gets brighter may provide a better contrast/lower background fluorescence for identifying regions of a neuronal circuit experiencing activity without the need for restricting expression to the soma.

## Imaging mouse hippocampal CA1 in brain slice at 5 kHz using variable light levels

Imaging neuronal activity in the CA1 region of the hippocampus was achieved by injection of an AAV coding for Ulla under the control of the human synapsin promoter (Figure 6A). Two weeks after injection, optical measurements from hippocampal brain slices during Schaffer collateral electrical stimulation were recorded (Figure 6B and 6C). Upon stimulation a rapid increase in fluorescence could be detected near the stimulating electrode corresponding to neurons experiencing a depolarization of the plasma membrane. The depolarization was quickly followed by a decrease in fluorescence corresponding to neuronal inhibition previously observed in the CA1 region of the hippocampus with the GEVI ArcLight [32, 33]. The excitatory signal (0.2% ΔF/F)was easily observed in a single trial due to a robust Signal-to-Noise ratio (SNR ∼20) (Figure 6C).

**Figure 6.**
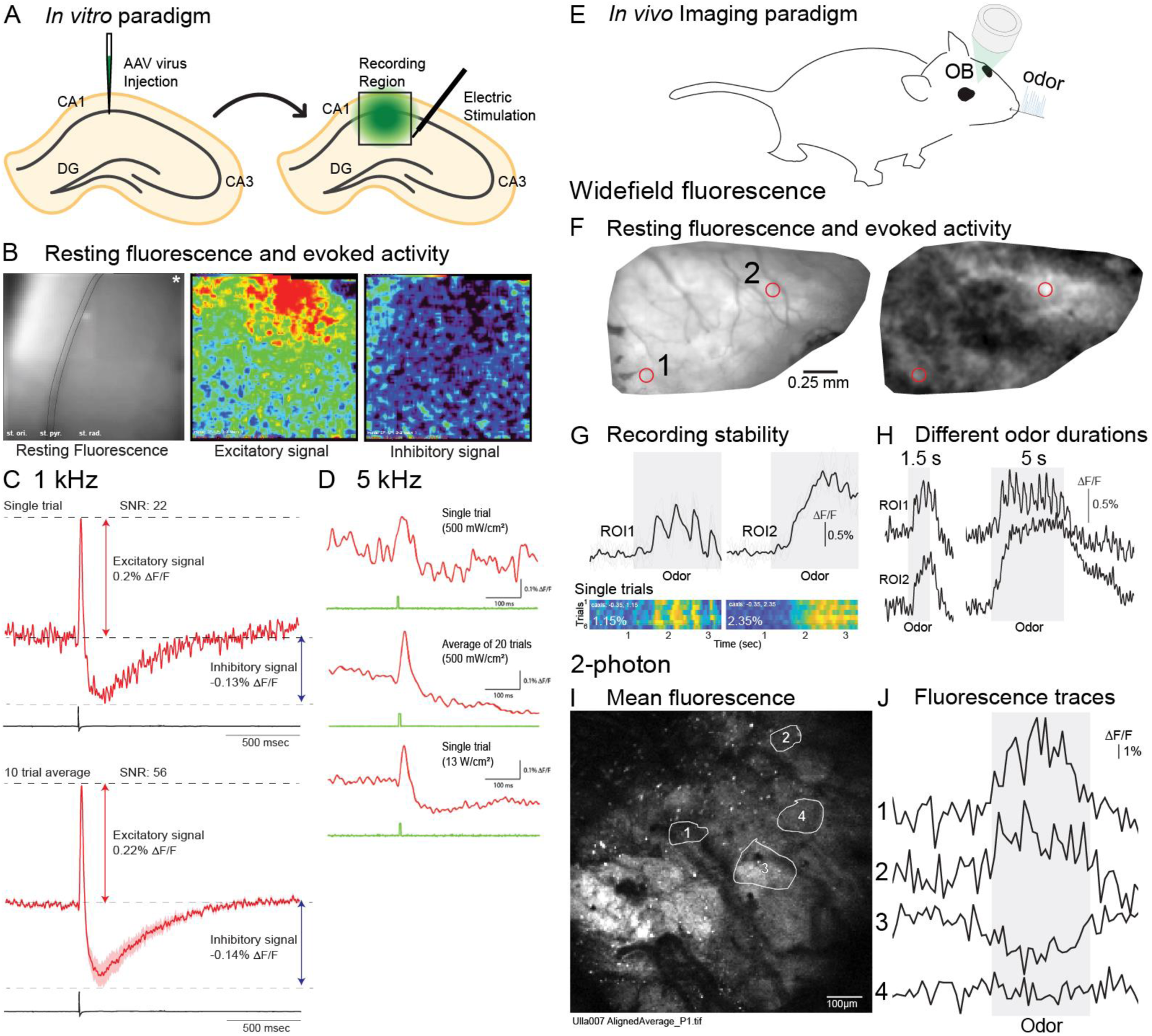
*Ex vivo* and *in vivo* population recordings using Ulla. **(A)** Schematic of acute brain slice recording from mouse hippocampal CA1 region at 1 kHz. Ulla was expressed in the CA1 region of the hippocampus. (**B**) Stimulation of Schaffer collaterals from CA3 (black trace in optical recording) yields a depolarization event followed by an inhibitory signal. Location of stimulation represented by an asterisk in the resting light image. **(C-D)** Acute brain slice recording at 1 kHz and 5 kHz under varying illumination intensities. Experimental design is the same as in **A**. Top trace is of the lowest light intensity for the LED (500 mW/ cm^2^). Averaging multiple trials improves the SNR as does increasing the illumination intensity (bottom trace). (**E**) *In vivo* imaging paradigm from the mouse olfactory bulb. (**F-H**) Epifluorescence imaging. (**F**) Mean in vivo fluorescence (left) and a frame subtraction analysis illustrating spatial patterns of odor evoked activity. (**G**) Mean (top) and single trial (bottom) fluorescence time course from two locations (red regions of interest in panel **F**). (**H**) Response from the same regions of interest in response to different odor stimulation durations. (**I**-**J**) 2-photon imaging. (**I**) Mean fluorescence from an average of 13 trials. (**J**) Fluorescence time course odor responses from four regions of interest in panel I. The recordings are from an average of 13 respiration aligned trials.

To investigate the illumination required, we imaged the CA1 region in brain slice again during electrical stimulation of the Schaffer collaterals at different excitation light levels and at a higher frame rate (5 kHz) to further reduce the number of photons detected per frame (Figure 6D). At the lowest setting of the LED (500 mW/cm^2^), a signal was measured in a single trial (0.2% ΔF/F, SNR 2), which was improved with averaging (Figure 6D, *top two traces*). Single trials with higher SNR were also observed at 5 kHz acquisition by increasing the power of the excitation light to 13 W/cm^2^.

## In vivo imaging of Ulla from the dorsal olfactory bulb using widefield fluorescence and 2-photon microscopy

To validate whether Ulla works *in vivo*, imaging was carried out from the olfactory bulb of mice that received an injection of an AAV expressing Ulla (Figure 6E). A diffuse fluorescence was evident in the olfactory bulb of all mice that received an AAV injection (Figure 6F, *left*). The fluorescence emission from the injected and uninjected hemibulbs were simultaneously measured using a low magnification lens. The injected side was 2.2 ± 0.5-fold brighter than the uninjected side (1.85-3.02 fold brighter, n = 4 preparations, p = 0.02 using a ranksum test). Thus, the AAV-serotype-promoter combination used in this study resulted in Ulla expression that was significantly brighter at baseline than endogenous autofluorescence present in uninjected hemibulbs. The odor methyl valerate was presented between 1-6% of saturated vapor evoked diffuse peaks of activity across the dorsal bulb in all tested preparations (Figure 6F, *right,* n = 9 preparations). Caudal and rostral regions of the olfactory bulb (ROIs 1 and 2) had distinct response dynamics. The caudal bulb exhibited distinct oscillations consistent with the animal’s respiratory pattern which tracked the duration of the odor presentation, while the rostral bulb was slower and had a more sustained response (Figure 6G, *left vs right*). The average response across all regions of interest in 3 preparations was 0.76 ± 0.28% ΔF/F (the response from the caudal and rostral ROI was measured in 4-5 trials for each preparation and averaged together; range: 0.38-1.07% ΔF/F). The recordings were relatively stable across individual trials within the same preparations, as illustrated by single trial measurements aligned to the odor onset with similar time course and amplitude signals (Figure 6G, *bottom heat maps*). Odors presented at 1.5 and 5 s evoked distinct numbers of respiratory oscillations illustrating that Ulla has sufficient temporal dynamics to track the known respiratory coupled oscillations present in the bulb (Figure 6H) [34-36]. Overall, the SNR and response dynamics were relatively stable across the single trials and comparable to those seen with ArcLight (Figure 6C, bottom heat maps) [37].Thus, we conclude that Ulla is a GEVI with sufficient brightness and SNR to detect odor-evoked signals *in vivo* using epifluorescence. However, since not all GEVIs work equally well under 2-photon excitation conditions, we next tested whether Ulla also could report odor evoked activity using 2-photon imaging. Following epifluorescence imaging to identify areas within the bulb activated by odor stimuli, mice were transferred to a 2-photon microscope for further analysis. Although the AAV expressing Ulla was not specific to one particular cell type, individual glomeruli could be more easily resolved using 2-photon excitation (Figure 6I). Odor-locked signals could be detected in localized regions within the olfactory bulb in respiration aligned trials (Figure 6J, *average of 13 trials*). Of the 4 animals imaged under 2-photon imaging, excitatory odor-locked signals were observed in all animals, inhibitory odor-locked signals were only observed in 1 animal. Therefore, Ulla can report odor-evoked activity in the mouse olfactory bulb using epifluorescence and 2-photon microscopy.

## Discussion

Genetically encoded fluorescent biosensors vary their fluorescence in response to changes in environmental conditions. By examining the fluorescence transitions of mutant GEVIs in conjunction with determining the structures of the FP domain in various pH conditions, we demonstrate that the mechanism of the voltage-dependent optical signal of ArcLight-derived GEVIs is distinct from the pH response and involves a conformational change inside the FP domain. In addition, we show that replacement of a bulky sidechain near the chromophore with threonine (F165T) introduces a distinct secondary component to the fluorescence transition likely due to increased flexibility of the chromophore. Remarkably, the orientation of the threonine sidechain at position 165 can be influenced by the composition of the flanking external amino acids creating a molecular switch to modulate the fluorescence of the biosensor.

The crystal structures of SE and SE A227D are virtually identical over a broad pH range indicating that the optical response to pH does not involve a conformational change (Supplemental figure S1). Rather, the pH response appears to be due to the sidechain of D147 forming a hydrogen bond with the chromophore while the other oxygen is exposed to the external solvent (Figures 1B and 1C). This creates a path from the external surface to the chromophore and was also seen in the structure of Lime which is a SE-derived FP [24]. Indeed, introduction of the S147D mutation in EGFP shifted the pKa to 7.2 enabling optical responses to changes in physiological pH range of the cell [26].

Despite creating a pathway from the surface to the chromophore, D147 does not appear to be directly involved in the voltage-dependent optical response. Introduction of the D147A mutation did not significantly alter the voltage response of the GEVI (Figure 2B) refuting the hypothesis that A227D commandeers the pH response of a neighboring domain via D147. Another pathway must exist for the voltage-dependent optical signal. That pathway can be influenced by the 147 position since D147K and D147N reduced the signal and introduced a secondary component to the fluorescent transition when membrane potential was altered (Figure 2B). Marina, an ArcLight-derived GEVI, which also has the D147A mutation inverted the polarity of the optical response to voltage [38] also suggesting an influence of position 147 on the voltage signal.

Another potential pathway for the voltage dependent optical signal involves the T203/T204 positions. Mutations to T204 exhibited a wide range of effects on the voltage-dependent optical signal including inverting the polarity of the signal when a negative charge was introduced (Figure 2D). The sidechain of the neighboring internal amino acid, T203, is located near the chromophore and can adopt different orientations in other FP structures (Figure 1C) [13]. Interestingly, T203 adopts only one orientation in SE and SE A227D regardless of pH. However, mutations to T203 exhibited altered kinetics for the fluorescence transition in response to voltage strongly implicating an internal conformational change as the cause of the voltage-dependent signal (Figure 3B).

The introduction of polar asymmetry to external neighboring amino acids in a β-sheet can bias the orientation of internal amino acid enabling threonine to become a molecular switch modulating the fluorescence depending on environmental conditions. The replacement of a bulky sidechain near the chromophore with threonine (F165T) created a fluorescence transition consisting of two distinct components (Figure 4A.). Bulky sidechains at 165 exhibit a simple fluorescence dimming in response to depolarization of the plasma membrane. Using the flanking amino acids of T203 in SE as a model, we introduced asymmetrical polarity with threonine and phenylalanine flanking the F165T mutant (Figure 4B). The directionality of that asymmetric polarity influenced the fluorescence transition indicating that the orientation of the internal threonine sidechain was influenced by the neighboring amino acids in the β-sheet.

The ability to influence the orientation of internal threonine sidechains was not restricted to the F165T mutant. The T204E/F223D mutations to the FP domain yielding the GEVI, Ulla, inverts the optical polarity of the voltage-induced transition (Figure 5A). Despite this inverted voltage response, the pH response of Ulla still gets brighter in alkaline conditions again revealing a distinction between the pH-dependent optical response and the voltage-dependent optical transition. The crystal structures of the Ulla FP domain revealed that T203 can adopt different orientations depending on the pH (Figure 5C). In addition the 7^th^ β-sheet is extended when the pH is above the pKa of Ulla flipping D147 to the external surface allowing H148 to interact with the chromophore.

The novel characteristics of Ulla, getting brighter during depolarization of the plasma membrane and the shift in the pH response, offer a relatively easy to use GEVI for resolving the activity neuronal circuits. Unlike calcium indicators, the expression of a fluorescent protein in the processes of cells can create a high background fluorescence [32]. A GEVI that starts dim and gets brighter may offer an improved contrast for detecting neuronal activity. For instance, Ulla was able to report neuronal activity in hippocampal slice under low light levels (Figure 6D). Ulla was also able to report neuronal activity *in vivo* under both 1-P and 2-P illumination from the mouse olfactory bulb (Figures 6F-J).

At two internal positions inside the FP domain, we were able to exploit the threonine sidechain as a molecular switch to modulate the fluorescence of the GEVI by altering the neighboring external chemistry. The first was at position T203 which has previously been proposed to function as a molecular switch based on crystal structures and molecular simulations [3, 13, 27, 39, 40]. The original SE introduced the external S202F and Q204T mutations resulting in a very rigid internal T203 sidechain that does not change orientation in response to pH. Introducing T204E inverted the voltage-dependent optical signal of the GEVI. The crystal structures of Ulla (T204E/F223D) revealed that T203 was oriented differently when the pH was below the pKa and that T203 did adopt different orientations in response to pH.

The second threonine switch was introduced by replacing a bulky sidechain near the chromophore, F165T. This threonine switch exhibited alternate fluorescence transition depending on the amino acid composition at position 164 and 166. The resting fluorescence and fluorescence transition can be controlled/influenced by the flanking external amino acid composition. Hence, these switches should be transferable to other FPs enabling the generation of multiple GEVIs with different optical spectra. In addition, these molecular switches may also aid the development of genetically encoded fluorescent biosensors ranging from ion detection to mechanosensors.

## Acknowledgements

In loving memory of Lawrence B. Cohen who provided critical reviews of the manuscript. We thank J. E. Dotzlaf and A. J. Kreuzman for their mentorship into protein structure and function. Molecular graphics and analyses performed with UCSF Chimera, developed by the Resource for Biocomputing, Visualization, and Informatics at the University of California, San Francisco, with support from NIH P41-GM103311. This study was funded by Korea Institute of Science and Technology grant 2E31523.

