## Supplemental material for "Exploiting threonine sidechains as molecular switches to modulate the fluorescence of genetically encoded biosensors"

**Supplemental data**

| Construct | State | Weighted τ (msec) | Fast τ (msec) | Slow τ (msec) | % fast |
|---|---|---|---|---|---|
| TM-SE | On | 9.5 ± 0.4 | 9.5 ± 0.5 | - | 100 |
| | Off | 10.5 ± 0.2 | 10.5 ± 0.2 | - | 100 |
| TM-SE T204E | On | 10.8 ± 0.4 | 8.9 ± 1.0 | 21.2 ± 1.2 | 85.7 ± 0.1 |
| | Off | 11.4 ± 0.6 | 6.0 ± 0.6 | 18.0 ± 0.2 | 55.1 ± 0.1 |
| TM-SE T204S | On | 8.6 ± 0.3 | 8.2 ± 0.4 | 20.1 ± 0.9 | 96.6 ± 2.2 |
| | Off | 10.2 ± 0.4 | 10.2 ± 0.4 | - | 100 |
| TM-SE T204N | On | 6.5 ± 0.5 | 6.5 ± 0.5 | - | 100 |
| | Off | 7.0 ± 0.3 | 7.0 ± 0.3 | - | 100 |

| Construct | State | Weighted τ (msec) | Fast τ (msec) | Slow τ (msec) | % fast |
|---|---|---|---|---|---|
| TM-SE T203N | On | 37.8 ± 2.6 | 9.1 ± 0.7 | 75.7 ± 5.8 | 56.6 ± 0.1 |
| | Off | 33.1 ± 2.2 | 11.1 ± 0.5 | 69.7 ± 6.2 | 62.0 ± 0.1 |
| TM-SE T203Q | On | 7.8 ± 0.6 | 7.8 ± 0.6 | - | 100 |
| | Off | 7.8 ± 0.3 | 7.2 ± 0.2 | 17.6 ± 1.1 | 94.3 ± 0.1 |
| TM-SE T203K | On | 36.9 ± 5.8 | 12.6 ± 1.0 | 74.0 ± 17.4 | 57.9 ± 3.5 |
| | Off | 28.8 ± 3.1 | 24.6 ± 2.3 | 64.0 ± 13.2 | 90.4 ± 9.6 |
| TM-SE T203S | On | 7.5 ± 0.6 | 7.5 ± 0.6 | - | 100 |
| | Off | 7.8 ± 0.4 | 7.8 ± 0.4 | - | 100 |

| Construct | State | Weighted τ (msec) | Fast τ (msec) | Slow τ (msec) | % fast |
|---|---|---|---|---|---|
| TM-SE T204E/F223D/A227 (Ulla) | On | 9.0 ± 0.3 | 9.0 ± 0.3 | - | 100 |
| | Off | 10.7 ± 0.4 | 10.7 ± 0.4 | - | 100 |

Supplementary table 2

| | SEA227 | | | | SEA227D | | | | SET204EF223DA227 | | | |
|---|---|---|---|---|---|---|---|---|---|---|---|---|
| | pH 4.5 | pH 5.5 | pH 7.5 | pH 9.5 | pH 4.5 | pH 5.5 | pH 8.0 | pH 9.5 | pH 4.5 | pH 6.0 | pH 7.0 | pH 8.5 |
| Data collection | | | | | | | | | | | | |
| Beam Line | PLS BL-5C | | | | | | | | | | | |
| Wavelength, Å | 0.97934 | | | | | | | | | | | |
| Space group | P212121 | P212121 | P212121 | P212121 | P212121 | P212121 | P1211 | P212121 | P212121 | P212121 | P1211 | P1211 |
| Unit cell parameters | | | | | | | | | | | | |
| a,b,c (Å) | 75.7, 80.6, 179.3 | 76.3, 79.9, 179.3 | 78.1, 78.3, 179.4 | 76.5, 80.3, 179.7 | 73.9, 82.9, 180.1 | 75.8, 80.5, 180.7 | 41.6, 53.4, 98.1 | 76.2 80.9 179.8 | 35.2, 46.1, 123.2 | 35.1, 46.4, 121.8 | 38.1, 122.8, 58.6 | 70.8, 48.2, 146.5 |
| α,β,γ (°) | 90=90=90 | 90=90=90 | 90=90=90 | 90=90=90 | 90=90=90 | 90=90=90 | α,γ=90, β=91.1 | 90=90=90 | 90=90=90 | 90=90=90 | α,γ=90, β=101.0 | α,γ=90, β=93.7 |
| aResolution, Å | | | | | | | | | | | | |
| No. of total reflections | 3102174 | 4394312 | 2607991 | 2699231 | 2423378 | 3341781 | 600725 | 822658 | 736615 | 810081 | 769233 | 1970348 |
| No. of unique reflections | 81052 | 56493 | 65254 | 112495 | 103630 | 25918 | 43183 | 31533 | 17891 | 36385 | 39278 | 56046 |
| No. in asymmetric unit | 4 | 4 | 4 | 4 | 4 | 4 | 2 | 4 | 1 | 1 | 2 | 4 |
| aCompleteness, % | 99.2 (96.3) | 97.4 (94.4) | 94.4 (80.3) | 97.7 (83.0) | 99.2 (98.3) | 95.1 (87.5) | 97.1 (89.0) | 97.4 (91.8) | 98.5 (96.3) | 98.7 (91.9) | 91.9 (65.2) | 84.4 (75.5) |
| aI/σ(I) | 13.5 (1.9) | 16.8 (3.1) | 16.5 (2.0) | 25.7 (3.3) | 16 (2.3) | 6.4 (1.8) | 20.0 (2.1) | 5.9 (1.4) | 16.6 (2.6) | 28.0 (2.8) | 19.0 (3.1) | 15.2 (3.0) |
| bRmerge, % | 12.6 (41.5) | 10.9 (39.7) | 12.2 (39.8) | 9.5 (43.4) | 14.0 (46.1) | 19.5 (32.1) | 8.6 (34.4) | 21.2 (45.0) | 10.1 (35.6) | 12.5 (39.4) | 7.9 (24.2) | 8.4 (23.4) |
| aCC1/2, % | 98.0 (28.0) | 99.3 (65.3) | 99.3 (39.0) | 99.7 (57.2) | 99.2 (62.8) | 93.4 (61.8) | 99.6 (73.0) | 93.6 (56.6) | 99.5 (45.1) | 98.8 (80.2) | 99.4 (88.0) | 99.1 (82.5) |
| Refinement | | | | | | | | | | | | |
| Resolution, Å | 50.0-1.95 (2.02-1.95) | 50.0-2.20 (2.28-2.20) | 50.0-2.10 (2.18-2.10) | 50.0-1.75 (1.81-1.75) | 50.0-1.80 (1.86-1.80) | 50.0-2.90 (3.00-2.90) | 50.0-1.76 (1.82-1.76) | 50.0-2.70 (2.80-2.70) | 50.0-1.85 (1.92-1.85) | 50.0-1.45 (1.50-1.45) | 50.0-1.95 (2.02-1.95) | 50.0-2.10 (2.18-2.10) |
| cRcryst/Rfree, % | 20.1/24.6 | 18.6/22.9 | 20.9/26.5 | 19.1/21.7 | 18.6/22.2 | 19.4/27.5 | 21.3/24.9 | 20.5/28.2 | 19.7/24.0 | 18.1/20.3 | 20.2/22.8 | 19.5/24.5 |
| No. of protein atoms | 7228 | 7228 | 7235 | 7231 | 7236 | 7267 | 3599 | 7263 | 1807 | 1804 | 3609 | 7211 |
| No. of water molecules | 575 | 523 | 341 | 816 | 883 | 0 | 263 | 0 | 138 | 246 | 259 | 385 |
| No. of ligand molecules | 4 | 4 | 4 | 4 | 4 | 4 | 2 | 4 | 1 | 1 | 2 | 4 |
| RMSD from ideal geometry: | | | | | | | | | | | | |
| Bond lengths, Å | 0.008 | 0.008 | 0.008 | 0.008 | 0.008 | 0.004 | 0.007 | 0.01 | 0.008 | 0.006 | 0.008 | 0.007 |
| Bond angles, º | 1.121 | 1.061 | 1.013 | 1.121 | 1.032 | 0.737 | 1.094 | 1.169 | 1.049 | 1.04 | 1.186 | 1.049 |
| Average B-factor, Å2 | 30.4 | 29.1 | 36.6 | 22.8 | 23.1 | 32.4 | 28.5 | 31.9 | 21.2 | 18.3 | 28.5 | 26.3 |

| | | | | | | | | | | | | |
|---|---|---|---|---|---|---|---|---|---|---|---|---|
| Ramachandran analysis | | | | | | | | | | | | |
| Favored, % | 95.9 | 95.3 | 95.7 | 96.4 | 96.4 | 93.9 | 96.6 | 93.8 | 97.7 | 96.8 | 96.8 | 96.3 |
| Allowed, % | 4.1 | 4.7 | 4.3 | 3.6 | 3.6 | 6 | 3.4 | 6.2 | 2.3 | 3.2 | 3.2 | 3.7 |
| Outliers | 0 | 0 | 0 | 0 | 0 | 0.1 | 0 | 0 | 0 | 0 | 0 | 0 |
| PDB code | 8JKC | 8JKG | 8JKI | 8JL2 | 8JL5 | 8JL6 | 8JL7 | 8JLL | 8JLM | 8JLS | 8JLT | 8JLU |

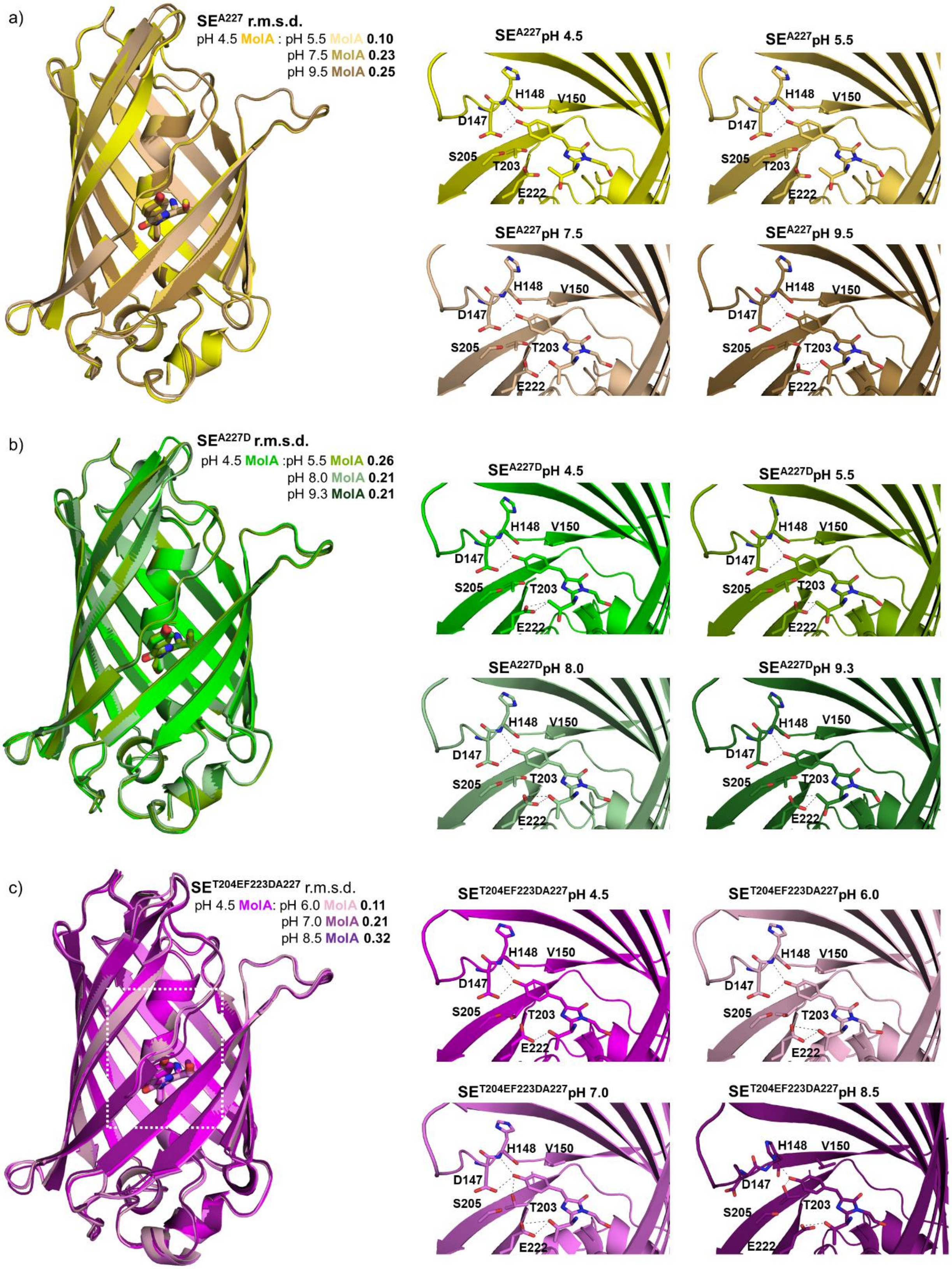
a)
SEA227 r.m.s.d.
pH 4.5 MolA : pH 5.5 MolA 0.10
pH 7.5 MolA 0.23
pH 9.5 MolA 0.25
SEA227pH 4.5
SEA227pH 5.5
SEA227pH 7.5
SEA227pH 9.5
H148
V150
D147
S205
T203
E222
b)
SEA227D r.m.s.d.
pH 4.5 MolA :pH 5.5 MolA 0.26
pH 8.0 MolA 0.21
pH 9.3 MolA 0.21
SEA227DpH 4.5
SEA227DpH 5.5
SEA227DpH 8.0
SEA227DpH 9.3
c)
SET204EF223DA227 r.m.s.d.
pH 4.5 MolA: pH 6.0 MolA 0.11
pH 7.0 MolA 0.21
pH 8.5 MolA 0.32
SET204EF223DA227pH 4.5
SET204EF223DA227pH 6.0
SET204EF223DA227pH 7.0
SET204EF223DA227pH 8.5

**Supplemental Figure 1. D147 provides a direct conduit from the exterior of the FP domain to the chromophore. A. The crystal structures of SE at pH 4.5, 5.5, 7.5, and 9.5.** Left is the overlay of the SE structures for all pH conditions. Right is the region near the chromophore for each pH condition. **B. The crystal structures of SE A227D at pH 4.5, 5.5, 8.0, and 9.3.** Left is the overlay of the SE A227D structures for all pH conditions. Right is the region near the chromophore for each pH condition. **C. The crystal structures of Ulla (SE T203E/F223D/A227 at pH 4.5, 6.0, 7.0, and 8.5.** Left is the overlay of the Ulla structures for all pH conditions. Right is the region near the chromophore for each pH condition. Potential hydrogen bonds are denoted by dashed lines. For all structures except Ulla pH 8.5, D147 forms a hydrogen bond with the chromophore. The other oxygen of the carboxyl side group of D147 is exposed to the solvent. For Ulla at pH 8.5 the 7th $\beta$-sheet begins at H148 which causes D147 to rotate entirely to the external surface. The T203 orientation for all structures of SE and SE A227D is consistent while T203 for Ulla is oriented differently and exhibits more flexibility as the pH increases. This is due to the mutation to the external F223 position (F223D).

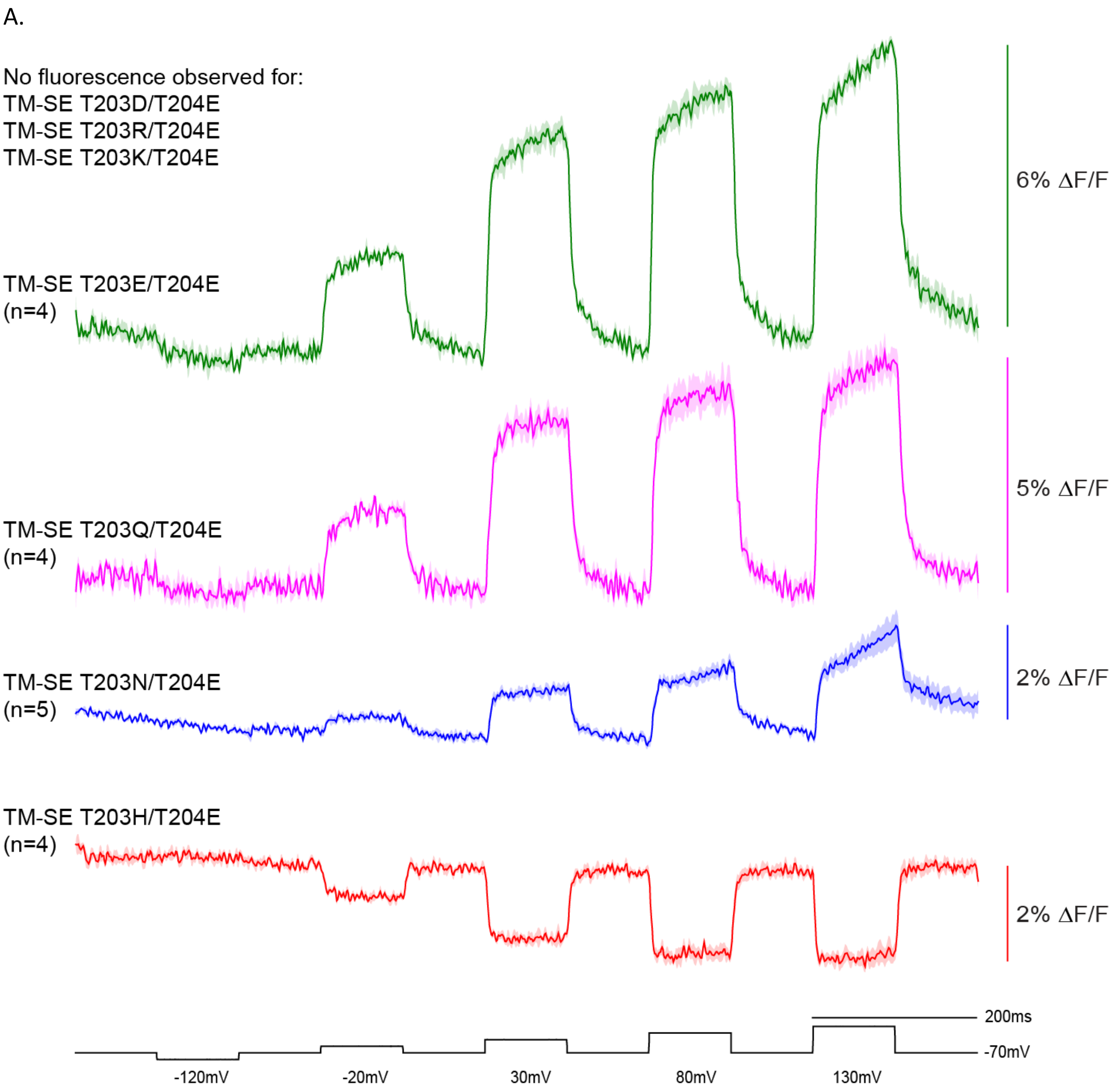
A.
No fluorescence observed for:
TM-SE T203D/T204E
TM-SE T203R/T204E
TM-SE T203K/T204E
TM-SE T203E/T204E
(n=4)
6% ΔF/F
TM-SE T203Q/T204E
(n=4)
5% ΔF/F
TM-SE T203N/T204E
(n=5)
2% ΔF/F
TM-SE T203H/T204E
(n=4)
2% ΔF/F
200ms
-70mV
-120mV
-20mV
30mV
80mV
130mV

B.

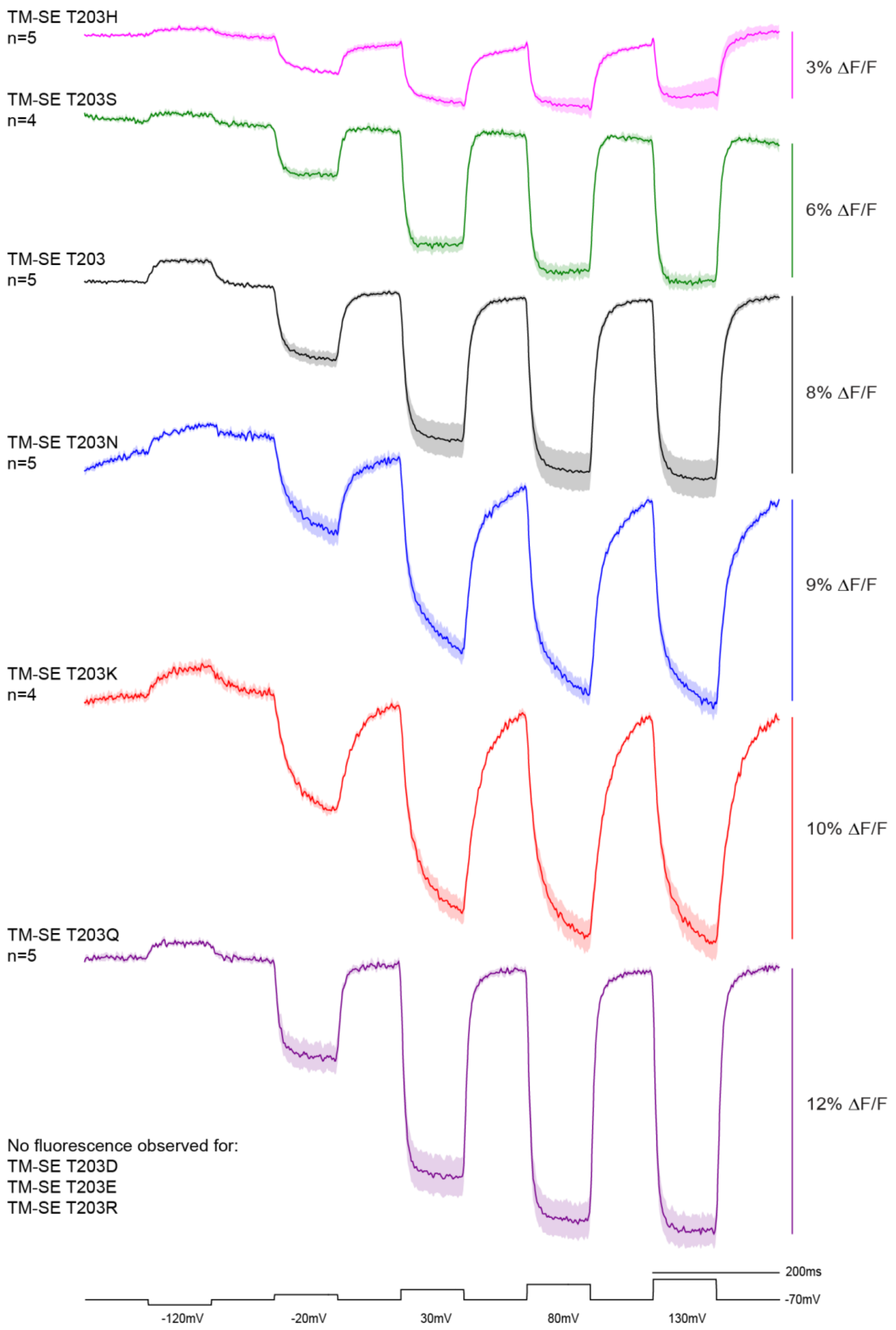
TM-SE T203H
n=5
3% ΔF/F
TM-SE T203S
n=4
6% ΔF/F
TM-SE T203
n=5
8% ΔF/F
TM-SE T203N
n=5
9% ΔF/F
TM-SE T203K
n=4
10% ΔF/F
TM-SE T203Q
n=5
12% ΔF/F
No fluorescence observed for:
TM-SE T203D
TM-SE T203E
TM-SE T203R
200ms
-70mV
-120mV
-20mV
30mV
80mV
130mV

**Supplemental figure 2. A. Double mutations to the T203 and T204 positions.** HEK cells expressing the indicated construct were subjected to whole-cell voltage clamp and imaged at 1Khz. **B. Single mutations to the internal T203 site affect the speed of the voltage-dependent optical signal.**
